# Replication initiation in bacteria: precision control based on protein counting

**DOI:** 10.1101/2023.05.26.542547

**Authors:** Haochen Fu, Fangzhou Xiao, Suckjoon Jun

## Abstract

Balanced biosynthesis is the hallmark of bacterial cell physiology, where the concentrations of stable proteins remain steady. However, this poses a conceptual challenge to modeling the cell-cycle and cell-size controls in bacteria, as prevailing concentration-based eukaryote models are not directly applicable. In this study, we revisit and significantly extend the initiator-titration model, proposed thirty years ago, and explain how bacteria precisely and robustly control replication initiation based on the mechanism of protein copy-number sensing. Using a mean-field approach, we first derive an analytical expression of the cell size at initiation based on three biological mechanistic control parameters for an extended initiator-titration model. We also study the stability of our model analytically and show that initiation can become unstable in multifork replication conditions. Using simulations, we further show that the presence of the conversion between active and inactive initiator protein forms significantly represses initiation instability. Importantly, the two-step Poisson process set by the initiator titration step results in significantly improved initiation synchrony with *CV* ∼ 1*/N* scaling rather than the standard 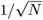 scaling in the Poisson process, where *N* is the total number of initiators required for initiation. Our results answer two long-standing questions in replication initiation: (1) Why do bacteria produce almost two orders of magnitude more DnaA, the master initiator proteins, than required for initiation? (2) Why does DnaA exist in active (DnaA-ATP) and inactive (DnaA-ADP) forms if only the active form is competent for initiation? The mechanism presented in this work provides a satisfying general solution to how the cell can achieve precision control without sensing protein concentrations, with broad implications from evolution to the design of synthetic cells.

## I. INTRODUCTION

Most biology textbooks explain biological decision-making by emphasizing the control and sensing of key protein concentrations through programmed gene expression and protein degradation in eukaryotes. Protein concentration gradients can encode spatial or temporal information across different scales, such as morphogen gradients in the French flag model in developmental biology [1] or cyclin oscillations in eukaryotic cell-cycle controls [2] [Fig. 1(a)]. However, in bacterial cell physiology, balanced biosynthesis has been the hallmark since the 1950s at the population and single-cell levels [3–5]. Balanced biosynthesis means that the synthesis rate of all cellular components is the same as the cell’s growth rate in steady-state growth, wherein the concentrations of stable proteins are steady by the balance of their production and dilution [Fig. 1(b)].

**FIG. 1.**
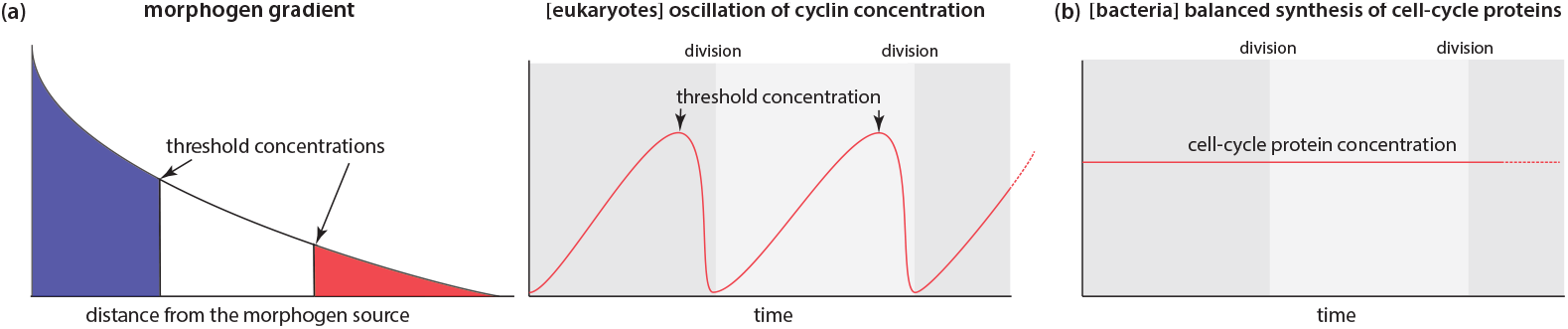
Protein concentration in eukaryotes vs. bacteria. (a) (left) morphogen gradient in the French flag model in developmental biology. (right) Oscillation of cyclin concentration for eukaryotic cell-cycle control. (b) Balanced biosynthesis in bacteria.

However, balanced biosynthesis poses a fundamental conceptual challenge to modeling the cell-cycle and cell-size controls, as the prevailing concentration-based models are not directly applicable if the concentration of cell-cycle proteins remains constant (within stochasticity). Indeed, for the billion-year divergent model bacterial organisms *Escherichia coli* and *Bacillus subtilis*, their size control is based on (1) balanced biosynthesis of division initiator protein FtsZ and (2) its accumulation to a threshold number (not concentration) [6]. These two conditions lead to the adder phenotype [6]. Unfortunately, a mechanistic investigation of threshold FtsZ number sensing is a formidable challenge because division initiation involves multiple interacting proteins with unknown properties [7].

Replication initiation in bacteria, which is exclusively controlled by the widely-conserved master regulator protein, DnaA, is an attractive problem for mechanistic investigation because it exhibits the adder phenotype [8– 11]. That is, the added cell size between two consecutive initiation events is independent of the cell size at initiation, as originally suggested by Sompayrac and Maaloe [12]. The adder phenotype implies that cells likely accumulate the DnaA molecules to a threshold number [6], and the synthesis of DnaA is balanced [13]. Furthermore, DnaA has been extensively studied, and most properties required for modeling are known or can be estimated [14–18]. Therefore, we view *E. coli* replication initiation as a tractable problem to understand the mechanism of protein copy-number sensing to control the cell cycle and cell size, and gain mechanistic insight into the general class of precision control in biology.

In this work, we revisit and significantly extend the initiator-titration model proposed by Hansen, Christensen, and Atlung thirty years ago [19], the model closest to the protein-number-sensing idea (see Section A). In Section A, we summarize the original initiator-titration model and introduce our initiator-titration model v2. In Section B, we first introduce the “protocell” model, a minimal version of the initiator-titration model, and derive the first expression of the protocell size at initiation (known as the “initiation mass”). In Section C, we perform a dynamical stability analysis of the protocell model and show the existence of initiation instability. In Section D, we extend the protocell to our “initiator-titration model v2” and derive an analytical expression for the initiation mass in a special case (the Δ4 mutant [13]) based on three mechanistic biological control parameters: the expression level of DnaA, the ratio of the active vs. passive forms of DnaA, namely, [DnaA-ATP]/[DnaA-ADP], and the number of DnaA titration boxes on the chromosome. In the same section, we show that adding the replication-dependent, biologically-observed DnaA-ATP →DnaA-ADP conversion element (RIDA) restores initiation stability [20, 21]. In Section E, we discuss initiation asynchrony and cell-to-cell variability using the concept of intrinsic and extrinsic noise in the framework of initiator-titration model v2.

Our model provides a quantitative and mechanistic explanation for several long-standing questions in bacterial replication initiation with the following findings: DnaA titration boxes are the protein-counting device that measures the threshold number of initiator proteins, and the two forms of DnaA (DnaA-ATP and DnaA-ADP), and especially the replication-dependent DnaA-ATP →DnaA-ADP, are needed to suppress initiation instability. Given the fundamental nature of replication initiation and its profound differences from eukaryotic cell-cycle control, we anticipate broad applications of our results, from the design of synthetic cells to the evolution of biological mechanisms in precision control.

## II. RESULTS AND DISCUSSION

### A. The “initiator-titration model v2” and intuition

Consider engineering a synthetic cell capable of self-replication. For such a cell to be viable, it must meet a fundamental requirement for cell-cycle control: initiating replication only once during cell division. A possible “simple” strategy to implement this requirement could be as follows [Fig. 2(a)]: (1) The chromosome has one origin of replication. (2) The cell produces one initiator protein during the division cycle. (3) The initiator protein binds to *ori* (the replication origin) and immediately triggers initiation. (4) Upon initiation, the cell destroys the initiator protein. While this seemingly straightforward strategy could limit the replication origin to a single site and produce a single initiator protein during cell division, the underlying mechanisms required to achieve this are likely more complex. For instance, how would the cell “know” when to produce them and when to degrade them?

**FIG. 2.**
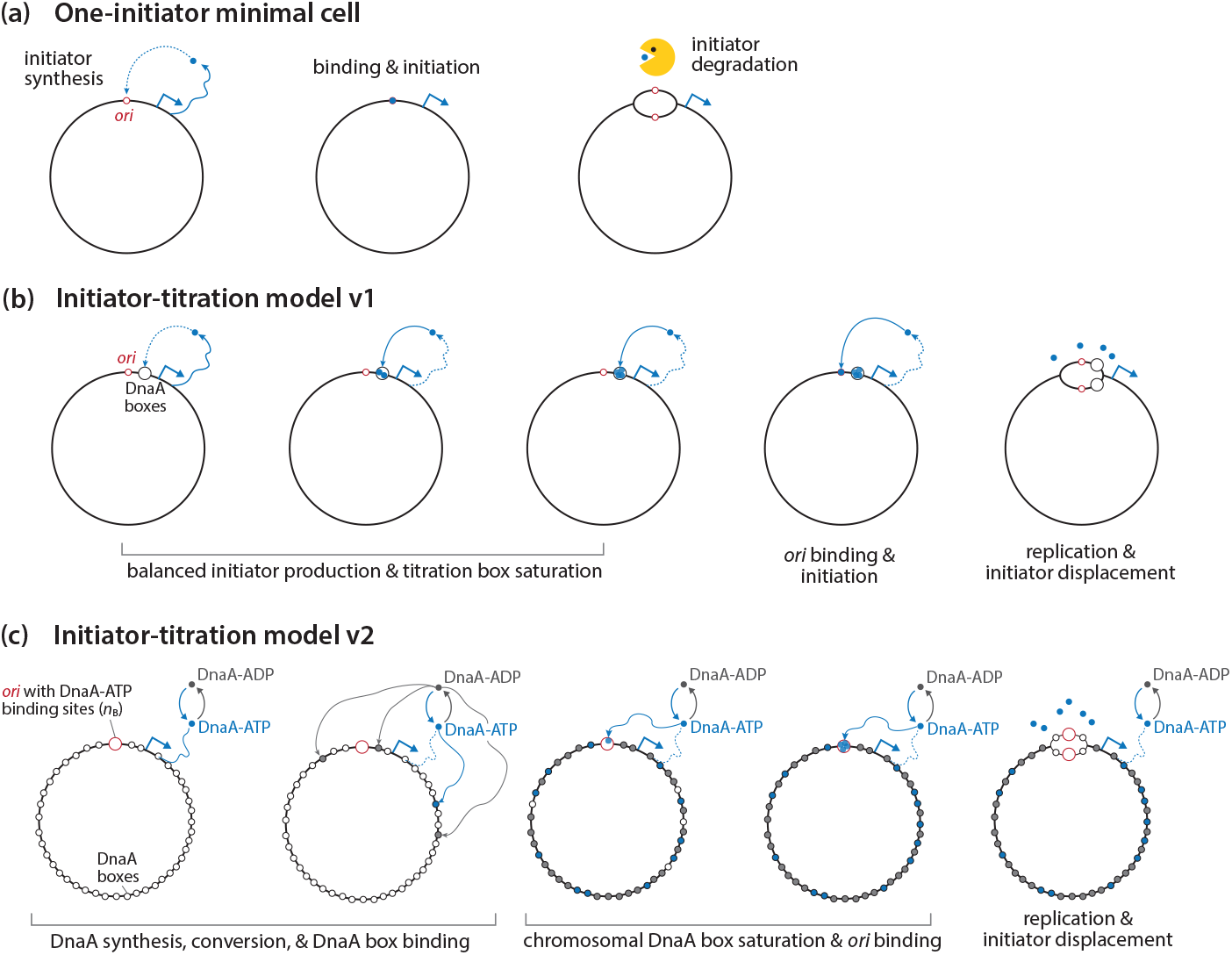
Initiation control. (a) Hypothetical minimal cell. (b) Initiator-titration model v1 [19]. (c) Initiator-titration model v2 (this work).

While *E. coli* exhibits characteristics similar to the hypothetical strategy described above, there are notable differences. *E. coli* has one replication origin (*ori*), but replication initiation requires 10-20 master regulator DnaA molecules binding to the eleven DnaA boxes at *ori* [15–17, 22]. Furthermore, DnaA is stable and not degraded upon initiation [15, 16]. Strikingly, *E. coli* produces approximately 300 copies of DnaA per *ori*, or 30 times more than required at *ori*, with almost all being titrated by DnaA boxes encoded on the chromosome [15, 16].

In 1991, Hansen and colleagues proposed the initiator-titration model to explain these observations [Fig. 2(b)] [19]. Their model posits that DnaA is first titrated by high-affinity DnaA boxes on the chromosome, which allows it to bind *ori* with weak affinity and initiate replication only after the chromosomal DnaA boxes are nearly saturated. This highlights the importance of DnaA boxes on the chromosome as the timing device for replication initiation.

Our model builds upon the initiator-titration model and incorporates the knowledge in DnaA accumulated in the past 30 years [15–17, 22]. Specifically, we have learned that DnaA exists in two forms, DnaA-ATP and DnaA-ADP, with different binding affinities to DNA [23]. DnaA-ATP is the active form that can trigger initiation, while DnaA-ADP is inactive as it cannot bind *ori* specifically [24, 25]. Further genetic, biochemical, and bioinformatic studies have revealed that approximately 300 high-affinity DnaA boxes are distributed across the circular chromosome [15, 26]. By contrast, *ori* contains a cluster of eleven DnaA binding sites, wherein only three have high affinities [25, 27]. Therefore, most DnaA, whether DnaA-ATP or DnaA-ADP, will first bind the high-affinity chromosomal DnaA boxes. Only after the titration step, do DnaA-ATP molecules bind the weak binding sites within *ori* and trigger initiation. We refer to this updated model as the initiator-titration model v2, in recognition of the pioneering work of Hansen *et al*. [19, 28].

Figure 2(c) illustrates how our initiator-titration model v2 works in more detail. To provide intuition without losing the generality of our ideas, let us consider a naked circular chromosome without bound DnaA.

1. As DnaA binds to ATP or ADP tightly [23] and the cellular concentration of ATP is almost 10x higher than ADP [29, 30], newly synthesized DnaA molecules become DnaA-ATP. During steady-state growth, both DnaA-ATP and DanA-ADP exist in the cell due to multiple interconversion mechanisms [16]. (See Section D and Appendix D for detailed discussion).
2. DnaA-ATP and DnaA-ADP will first bind to around 300 high binding-affinity chromosomal DnaA boxes (*K*_D_ ≈1 nM) [26], whereas only DnaA-ATP can bind to around ten low-affinity boxes within *ori* (*K*_D_ ≈ 100 nM) [26] [31].
3. When most chromosomal DnaA boxes are saturated, the probabilities for DnaA-ATP binding to *ori* vs. the remaining chromosomal DnaA boxes become comparable. Initiation is triggered once the low-affinity *ori* binding sites are saturated by DnaA-ATP.

As we elaborate below, the initiator-titration model v2 answers two long-standing fundamental questions:

1. Why does *E. coli* produce so many more DnaA proteins than required for initiation, only to be titrated?
2. Why does *E. coli* maintain two forms of DnaA in the first place if they only need DnaA-ATP for initiation?

### B. The “protocell”: a minimal initiator-titration model

To gain analytical insight, we first construct a minimal initiator-titration model, named “protocell” [Fig. 3(a)]. The protocell has the complexity between the two versions of the initiator-titration model [Figs. 2(b) & 2(c)]. The protocell has one *ori*, the active initiator protein (e.g., DnaA-ATP in *E. coli*), and the initiator binding sites on the chromosome. We assume the following based on the experimental data:

**FIG. 3.**
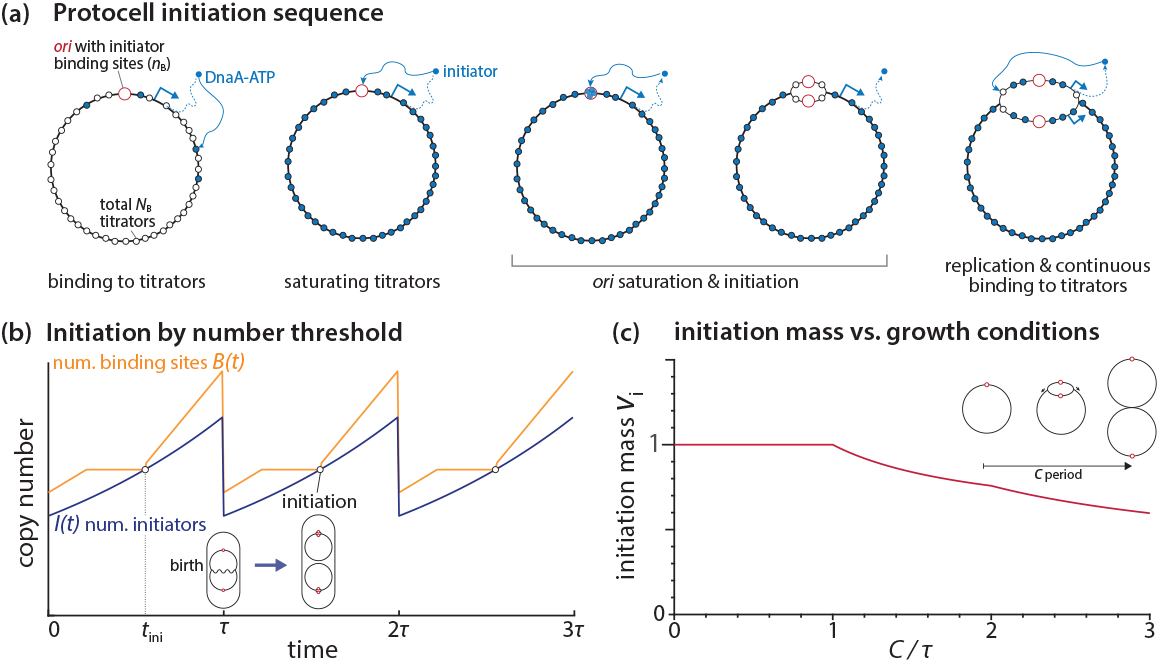
Initiation control of the protocell by initiator protein counting. (a) Model sequence of titration and initiation. (b) Change in the copy numbers of initiators and initiator binding sites during the cell cycle under the condition of two overlapping cell cycle (*C < τ < C* +*D*). The initiation condition is *I*(*t* = *t*_ini_) = *B*(*t* = *t*_ini_). (c) Predicted initiation mass in different growth conditions (*C/τ*) by assuming that *c*_I_ is a constant [32, 33].

1. The cell grows exponentially *V* (*t*) = *V*_0_*e*^*λt*^ in steady-state [3], where *V* (*t*) is the total cell size at time *t*, and *λ* is the growth rate. The mass-doubling time *τ* is given by 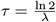.
2. Synthesis of the initiator protein is balanced, i.e., its concentration is constant during growth [3]. We denote the initiator protein copy number at time *t* as *I*(*t*) and its concentration as *c*_I_.
3. The rate of DNA synthesis is constant [34, 35], with the duration of chromosome replication *C*, independent of the mass-doubling time *τ* [36].
4. The chromosome encodes specific DNA sequences for binding of the initiator proteins. *N*_B_ high-affinity sites are evenly distributed on the chromosome [15], and *n*_B_ low-affinity sites are localized at *ori* [16]. For the *E. coli* chromosome, we set *N*_B_ = 300 and *n*_B_ = 10, as explained in Section A. During replication, the total number of initiator binding sites increases as *B*(*t*).
5. Initiators tightly bind to the binding sites rather than staying in the cytoplasm, and initiators preferentially bind the chromosomal binding sites before binding to the ones at *ori*. Therefore, replication initiates at *t* = *t*_ini_ when *I*(*t* = *t*_ini_) = *B*(*t* = *t*_ini_), namely, all binding sites are saturated by the initiator proteins.

For illustration purposes, we consider an intermediate growth condition, where two cell cycles slightly overlap without exhibiting multifork replication [36] [Fig. 3(b)]. In the Helmstetter-Cooper model [37], this corresponds to *C < τ < C* + *D*, where *D* is the duration between replication termination and cell division. As such, the cell can have two intact chromosomes between termination and the next initiation [Fig. 3(b)].

The steady-state curves of *I*(*t*) and *B*(*t*) are shown in Fig. 3(b) (in our model, a steady state means all derived quantities are periodic with a period of *τ*). In general, *I*(*t*) increases exponentially because of exponential growth and balanced biosynthesis (Assumptions 1&2 above), whereas *B*(*t*) increases piecewise linearly because of replication initiation and termination (Assumptions 3&4). Therefore, the number of initiators catches up with the total number of binding sites between replication termination and the new round of initiation at *I*(*t* = *t*_ini_) = *B*(*t* = *t*_ini_) = 2(*N*_B_ + *n*_B_) (Assumption 5). Here, the factor “2” refers to the fact that there are two entire chromosomes and two *ori*’s right before the initiation event in the specific growth condition depicted in Fig. 3(b). Upon initiation, the number of binding sites *B*(*t*) increases discontinuously by 2*n*_B_ due to the duplication of both *ori*’s and the binding sites therein. After that, *B*(*t*) increases at the rate 2*N*_B_*/C*, steeper than the slope of *I*(*t*). Once the cell divides, *I*(*t*) and *B*(*t*) drop by half, and the cell repeats its cycle.

From this picture, the initiation mass *v*_i_, defined by cell volume per *ori* at initiation [36], can be easily calculated by the number of initiators at initiation,

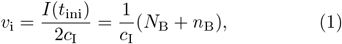

where *c*_I_ is the initiator protein concentration, and “2” reflects the copy number of *ori* before initiation.

The above result can be extended to different growth conditions. For example, in slow growth (*τ* > *C* + *D*), the replication cycles do not overlap, and all the factor “2” will vanish in the above analysis due to the single chromosome at initiation. This results in the same initiation mass *v*_i_ as in the intermediate growth condition. In fast-growth conditions (*τ* < *C*), replication cycles overlap, exhibiting multifork replication. Since a new round of replication starts before the previous round of replication is completed, the initiation mass is given by

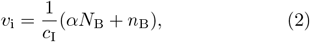

with the cell-cycle dependent parameter *α* ≤ 1 given as

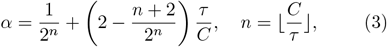

which applies to any growth conditions (see Appendix A for a derivation). *α* = 1 when *τ ≥C* (non-multifork replication), and 0 *< α <* 1 when *τ < C* (multifork replication) [Fig. 3(c)]. Thus, *α* refers to the degree of overlapping replication. Some of the most salient predictions of these results include (1) The initiation mass is inversely proportional to the initiator concentration *c*_I_(2) The initiation mass linearly depends on the number of chromosomal binding sites *N*_B_.

The basis of the protocell’s behavior is that the initiator increases exponentially, whereas the number of binding sites increases piecewise linearly only during DNA replication. This allows the cell to reach the initiation point *I*(*t*) = *B*(*t*) from any initial conditions. Therefore, the protocell can always trigger initiation by protein number counting through titration.

### C. The protocell exhibits initiation instability

In the last section, we addressed if a solution exists in the minimal protocell model with a period of *τ*. We showed that this periodic solution always exists (Eq. 2). We defined it as the “steady-state” solution in the biological sense that the cell can grow in a steady state with the periodic cell cycle. However, since the model is dynamic, convergence to a steady state from a given initial condition, *I*(0) and *B*(0), is not guaranteed. Hence, in this section, we study how the replication cycle propagates in the lineage from an arbitrary initial condition at *t* = 0, and under what conditions the cycle converges to the steady-state solution.

Intuitively, if the two consecutive initiations are separated by *τ*, thus periodic, the system is in a steady state.

5Suppose an initiation event at *t* = 0, and its initiation mass deviates from the steady-state solution Eq. 2. Typically, the next initiation occurs at *t* = *t*^+^ ≠ *τ*. However, if this time interval between two consecutive initiations eventually converges to *τ* after generations, the steady-state solution is stable under perturbations on the initial conditions. Otherwise, the steady-state solution is unstable.

In the rest of this section, we analyze a dynamical system based on Assumptions 1-5 in Section B on the protocell.

#### 1. Setup

We consider a protocell containing one chromosome with ongoing multifork replication [Fig. 4(a)]. We block the cell division so the protocell grows indefinitely as the chromosome replicates and multiplies starting from the initial condition. As the cell size approaches infinity, does the initiation mass have a fixed value (stable) or multiple values (unstable)? The analysis is non-trivial, as we need to accommodate arbitrary initial conditions.

**FIG. 4.**
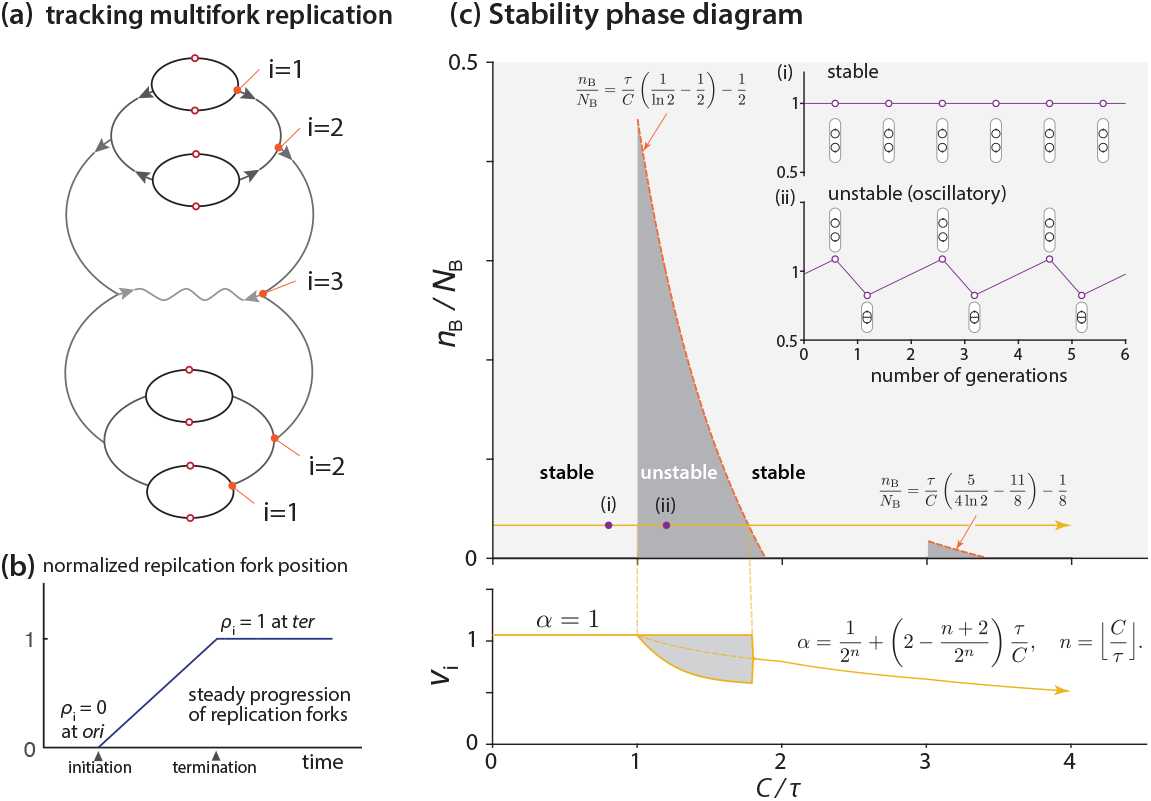
Dynamical stability analysis for initiation of the protocell. (a) Multifork replication tracker. *i* = 1 represents the group of replication forks closest to *ori*. (b) Linear (in time) progression of the replication forks from *ori* to *ter* on the circular chromosome. *ter* is on the opposite end of *ori* on the chromosome. (c) Stability phase diagram (*n*_B_*/N*_B_ vs. *C/τ* space). Below: initiation mass vs. *C/τ* in the condition of *n*_B_*/N*_B_ = 1*/*30 and constant *c*_I_ [32, 33]. Inset: stable initiation events vs. unstable (oscillatory) initiation events.

To this end, we start with the dynamics of *I*(*t*) and *B*(*t*). First, we have

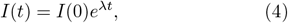

as a consequence of exponential cell growth and balanced biosynthesis of the initiator proteins. Next, the dynamics of the number of binding sites *B*(*t*) is more subtle because it increases piecewise linearly depending on the replication state of the chromosome and the number of replication forks. To accommodate the possibility of arbitrary initial conditions, we define the “multifork tracker” vector variable, ***ρ***(*t*), as follows.

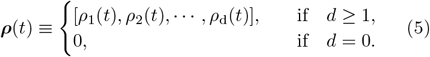

Here, the index *d* is the total number of generations (namely, the total rounds of replication cycles) since the initial chromosome, so *d* can grow indefinitely with time. That is, at every new round of the replication cycle, the size of the vector increases by one from *d* to *d* + 1. *d* = 0 is for the initial cell supposed to have an intact single chromosome without ongoing replications.

We use the variable *ρ* to indicate the relative position of a replication fork of interest between *ori* and *ter* (the replication terminus), and therefore 0≤ *ρ*(*t*) ≤ 1 [Fig. 4(b)]. For example, *ρ* would be 0.5 if a pair of forks is exactly halfway between *ori* and *ter* [Figs 4(a) & 4(b)]. To track multifork replication, we use *ρ*_i_(*t*) to represent the group of replication forks that are the i-th closest to the *ori* [Fig. 4(a)]. For example, *i* = 1 always refers to the newest group of replication forks. To record the replication history, we set *ρ*_i_(*t*) = 1 for those replication forks that have already reached *ter* [Fig. 4(b)]. By these definitions, ***ρ***(*t*) applies to both multifork replication and non-multifork replication.

Based on the multifork tracker vector, the number of binding sites *B*(*t*) is completely determined by ***ρ*** as

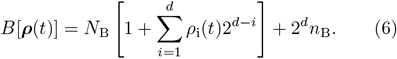

The dynamics of ***ρ***(*t*) consists of two parts: First, between two initiation events, *ρ*_i_(*t*) increases linearly with a slope of 1*/C* until it reaches 1, as replication forks travel from *ori* to *ter* [Fig. 4(b)]. Second, at initiation, the dimension of ***ρ*** increases by one, shifting its components to the right as *S* : ℝ^*d*^ → ℝ^*d*+1^, (*ρ*_1_, *ρ*_2_,*…, ρ*_*d*_)↦(0, *ρ*_1_, *ρ*_2_,*…, ρ*_*d*_) to accommodate the new pair of replication forks at each *ori* [see also, Fig. 4(a)].

#### 2. Properties of the steady state

The steady-state solution assumes periodicity of dynamics so that *I*(*t*) and *B*(*t*) double in each replication cycle. We consider the mapping between two consecutive initiation events to solve for the steady-state condition. We denote the first initiation event as ***ρ***(*t* = 0) = ***ρ*** at *t* = 0, and the second initiation event as ***ρ***(*t* = *t*^+^) = ***ρ*** ^+^ at *t* = *t*^+^. The mapping ℱ : ℝ^*d-*1^→ℝ^*d*^, ***ρ ↦ρ***^+^ requires a time-translation and a shift:

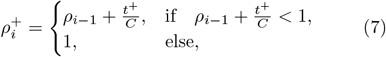

where the initiation time *t*^+^ is determined by the initiation criteria that *I*(*t* = 0) = *B*(*t* = 0) and *I*(*t* = *t*^+^) = *B*(*t* = *t*^+^), Eq. 4, and Eq. 6,

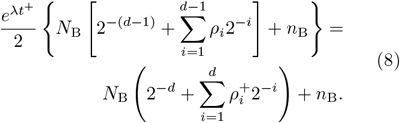

Equations 7 & 8 describe the dynamics of the system at initiation. We can now obtain the fixed point of the mapping ℱ by setting *d*→ *∞* and ***ρ***^+^ = ***ρ*** (Appendix B):

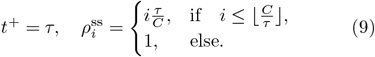

The resulting expression for steady-state initiation mass is the same as Eq. 2, i.e., the fixed point of ℱ is the steady-state solution (see Appendix B for more details).

Next, we study the stability of the fixed point of ℱ by calculating the Jacobian matrix of ℱ at the fixed point:

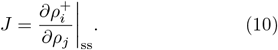

This matrix can be reduced to an *n*×*n* matrix 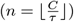, since all other matrix elements are zero. In *E. coli*, 0≤ *n* ≤2 in most growth conditions; here, we consider the range of 0≤*n*≤3 to accommodate cells theoretically doubling as frequently as at every *τ* = 10 minutes, with a C period of 40 minutes. Therefore, we can calculate the eigenvalues of *J* for each *n*. Stability requires the largest eigenvalue of *J* to be smaller than 1. Eventually, we can obtain the stable and unstable regimes in the *n*_B_*/N*_B_ vs. *C/τ* phase diagram, as shown in Fig. 4(c) (see Appendix B, also Appendix Fig. 2). Importantly, the phase diagram reveals both stable (*n <* 1) and unstable (small *n*_B_*/N*_B_ when *n >* 1) steady states [Fig. 4(c)].

What happens when the system becomes unstable? As discussed earlier, in fast growth conditions, *α <* 1 in the steady-state initiation mass expression (Eq. 2). Indeed, using numerical simulations, we found that the initiation mass oscillates between two values [Fig. 4(c)]. This indicates that the cell cycle can oscillate between multifork and non-multifork replication. Mathematically, this oscillatory behavior means that the fixed points of ℱ^*o*2^ = ℱ ◦ ℱ are stable, although the fixed point of ℱ is unstable. By fixing one of the fixed points of ℱ^*o*2^ as *ρ*_1_ = 1, we can compute the other fixed point with *ρ*_1_ *<* 1 (see Appendix C). In extreme cases, *ρ*_1_ can be as small as 0.1. That is, the second round of replication starts only after 10% of the chromosome has been replicated by the replication forks from the previous initiation. When the replication forks from two consecutive rounds of initiation are too close to each other, they cannot be separated into two division cycles. This should result in two initiation events in one division cycle, and no initiation in the next division cycle.

Therefore, although initiation triggering is guaranteed, the performance of the protocell is imperfect in terms of initiation instability in certain growth conditions. We show how the initiator-tiration model v2 resolves the instability issue in Section D.

### D. The initiator-titration model v2: replication-dependent DnaA-ATP→DnaA-ADP conversion stabilizes the cell cycle

In the previous section, we showed that the protocell can show initiation instability. In understanding why wild-type *E. coli* initiation is stable, we have to consider unique features of DnaA in *E. coli*, namely, its two distinct forms: the active DnaA-ATP and the inactive DnaA-ADP [23]. Several extrinsic elements, categorized into two main groups, interconvert between these DnaA forms [16, 20, 21, 38–41] [Fig. 5(a)].

**FIG. 5.**
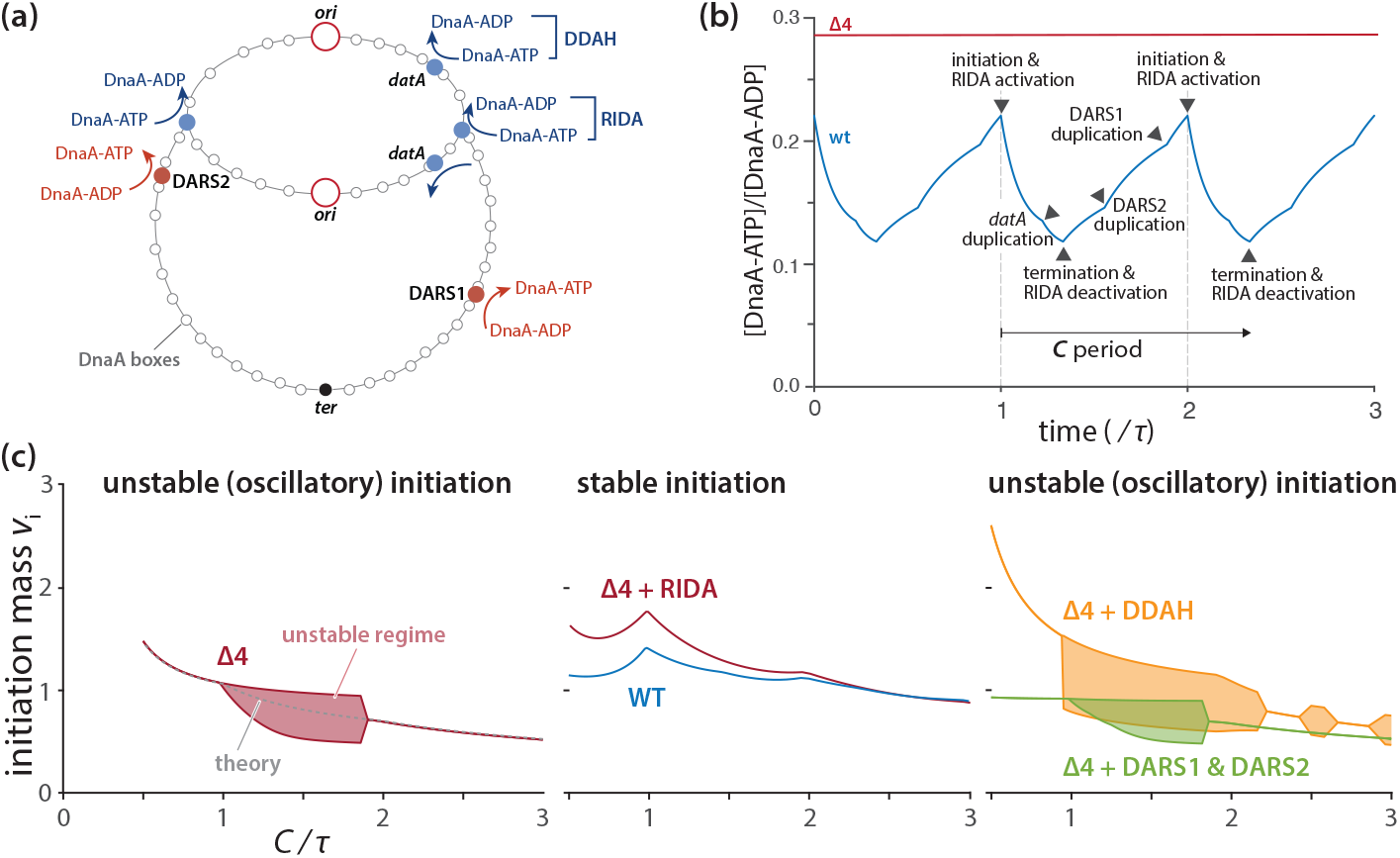
Initiator-titration model v2 predictions (see Section D for more details). (a) External DnaA-ATP *↔* DnaA-ADP conversion elements in *E. coli*. RIDA is the only component that depends on the active replication forks. (b) [DnaA-ATP]/[DnaA-ADP] varies during the cell cycle predicted by computer simulations in the wildtype cells. By contrast, the Δ4 mutant that lacks all extrinsic DnaA-ATP *↔*DnaA-ADP conversion elements in (a) show constant ratio. (c) Predicted initiation mass in different growth conditions (*C/τ* with fixed *C*) by assuming a constant [DnaA] [32, 33]. RIDA, the replication-dependent DnaA-ATP *→*DnaA-ADP mechanism alone can restore stability as long as titration is present. None of the other extrinsic DnaA-ATP *↔* DnaA-ADP conversion elements can restore the initiation stability.

The first group catalyzes the conversion of DnaA-ATP →DnaA-ADP. This includes the <u>R</u>egulatory Inactivation of <u>D</u>na<u>A</u> (RIDA) [20, 21] and *<u>d</u>atA*-dependent <u>D</u>naA-<u>A</u>TP <u>H</u>ydrolysis (DDAH) [38, 39]. RIDA’s functionality requires active replication forks [42], thus rendering it replication-dependent, while DDAH’s *datA*, a DnaA binding chromosomal locus, fosters DnaA-ATP hydrolysis.

By contrast, the second group, comprised of DARS1 and DARS2 (types of <u>D</u>na<u>A R</u>eactivating <u>S</u>equences), facilitates the ADP→ATP exchange for DnaA-ADP [40, 41].

Importantly, DnaA harbors intrinsic ATPase activity that facilitates its own conversion from DnaA-ATP →DnaA-ADP [23, 27], a feature not depicted in Fig. 5(a). Intriguingly, Δ4 cells — cells with a full deletion of all extrinsic DnaA-ATP ↔ DnaA-ADP interconversion pathways — exhibit a nearly identical initiation phenotype as wild-type cells [13], solely relying on DnaA’s intrinsic ATPase activity.

Figure 5(b) presents our numerical simulation results illuminating the alterations in the [DnaA-ATP]/[DnaA-ADP] ratio during the cell cycle in a DNA replication-dependent manner (see Appendix E for simulation details). Notably, this ratio should remain steady in Δ4 cells during cell elongation in a steady state [13]. Furthermore, re-initiation is prohibited within a certain timeframe post-initiation (approximately 10 minutes; the “eclipse period”), attributable to the sequestration of newly synthesized DNA by SeqA [43].

In this section, we integrate each of these features into our protocell model to formulate our initiator-titration model v2 and compute the initiation stability phase diagram. Our findings reveal that the replication-dependent DnaA-ATP→DnaA-ADP conversion by RIDA largely alleviates initiation instability, thus reinstating the stability characteristic of wild-type cells.

#### 1. Analytical expression of the initiation mass in the initiator-titration model v2 with a constant DnaA-ATP/DnaA-ADP ratio (Δ4 cells)

First, we incorporate the two forms of DnaA with the intrinsic DnaA-ATP →DnaA-ADP activity by DnaA into the protocell model to construct the Δ4 cells, a minimal version of the initiator-titration model v2 [Fig. 2(c)]. As noted earlier, both DnaA-ATP and DnaA-ADP can bind the chromosomal DnaA boxes because of their strong binding affinity (*K*_D_∼ 1 nM [15, 26, 44]), whereas only DnaA-ATP can bind the weak DnaA boxes at *ori* with *K*_D_ ∼ 10^2^ nM [15, 26, 45]. With the same Assumptions 1-4 in Section B and this additional assumption, we can derive an analytical expression for steady-state initiation mass for Δ4 *E. coli* (see Appendix D for the derivation):

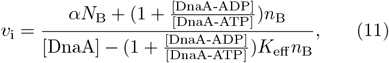

where *K*_eff_ is the effective dissociation constant of DnaA at *ori*, and *α* is in Eq. 3. Therefore, this equation brings together the expression level of DnaA via [DnaA], the ratio [DnaA-ATP]/[DnaA-ADP], and the degree of overlapping replication (*α*).

Note that, under physiological conditions, *K*_eff_ *n*_B_≪ [DnaA] (Appendix D). If [DnaA-ATP] ≫[DnaA-ADP], all DnaA molecules are in their active form DnaA-ATP, and the Δ4 *E. coli* converges to the protocell (i.e., Eq. 11 converging to Eq. 2).

#### 2. *The* Δ4 *cells show initiation instability*

We also investigated the initiation stability of the Δ4 cells using numerical simulations [Fig. 5(c)] (see Appendix E for simulation details). The initiation stability phase diagram is analogous to that of the protocells in Fig. 4(c), showing an island of instability regime. This occurs during the transition into multifork replication, wherein initiation mass alternates between two values. Importantly, changing the [DnaA-ATP]/[DnaA-ADP] ratio does not significantly impact the stability [Appendix Fig. 2(b)].

#### 3. Replication-dependent DnaA-ATP →DnaA-ADP by RIDA alone can restore initiation stability

Next, we implemented the extrinsic DnaA-ATP↔ DnaA-ADP conversion elements in the Δ4 cells. In contrast to the constant [DnaA-ATP]/[DnaA-ADP] in Δ4, the extrinsic conversion elements induce temporal modulations in [DnaA-ATP]/[DnaA-ADP] during cell elongation [46]. This ratio reaches its maximum at initiation and its minimum at termination due to the activation/deactivation of the RIDA mechanism [Fig. 5(b)] [47].

We also investigated the initiation stability across growth conditions [Fig. 5(c) and Appendix Fig. 3]. Among all the known extrinsic conversion elements we tested, the replication-dependent DnaA-ATP →DnaA-ADP by RIDA alone was sufficient to restore initiation stability [Fig. 5(c), also Appendix Fig. 3]. Other elements only had mild effects on the stability. RIDA is replication-dependent; thus, it immediately decreases the level of DnaA-ATP upon initiation. This reduction in the initiation-competent DnaA-ATP level is likely the reason for suppressing premature re-initiation.

Although we found RIDA to be the initiation stabilizer, it still significantly delays initiation due to the reduced level of DnaA-ATP. Our simulations show that the delayed initiation can be alleviated by the other DnaA-ADP→DnaA-ATP conversion elements without causing instability [Fig. 5(c) and Appendix Fig. 3]. Interestingly, the initiation mass becomes nearly invariant across a wide range of growth conditions in the presence of all four extrinsic conversion elements [Fig. 5(c)], as long as the concentration [DnaA] is growth-condition independent. We previously used this growth-condition-independent [DnaA] hypothesis to explain the invariance of initiation mass [36], and the data so far supports the hypothesis [32, 33].

Based on these results, we conclude that the replication-dependent DnaA-ATP→DnaA-ADP by RIDA can significantly enhance the initiation stability, and the other DnaA-ADP→DnaA-ATP conversion elements keep the initiation mass nearly constant against physiological perturbations.

#### 4. The eclipse period or origin sequestration does not improve stability

We also tested the effect of the eclipse period [43] in our simulations (Appendix Fig. 4). During the predefined eclipse period, we did not allow the binding of the initiator to *ori*. Surprisingly, the eclipse period did not improve stability significantly in the multifork replication regime. However, the amplitude of the initiation mass oscillation decreased slightly (Appendix Fig. 4). Therefore, we predict the effect of SeqA on steady-state stabilization to be modest.

#### 5. Comparison with previous modeling by Berger and ten Wolde and recent experimental work

In their recent study, Berger and ten Wolde [11] conducted a thorough investigation into *E. coli* DNA replication. They utilized extensive numerical simulations that factored in the known dynamics between DnaA-ATP and DnaA-ADP conversion, as well as the aspects of DnaA titration. To our knowledge, Berger and ten Wolde were the first to suggest possible instability during multifork replication.

Under relatively fast growth conditions (with the doubling time 35 mins and the C period 40 mins), their observations noted oscillations in the initiation mass between two distinct values, which occurred in the absence of DnaA-ATP↔DnaA-ADP conversion. Our instability phase diagram [Fig. 4(c)] explains this observation. For example, in the case of Δ4 mutant cells, the initiation mass should oscillate between two values when 1 *< C/τ <* 1.8 [Fig. 5(c)]. However, the complexity of these instability regimes needs to be noted. Our phase diagrams show that multifork replication does not always lead to instability [Fig. 5(c) and Appendix Fig. 2].

Berger and ten Wolde propose the DnaA-ATP↔ DnaA-ADP conversion as the key mechanisms in initiation control, as DnaA-ATP↔ DnaA-ADP conversion could avoid initiation instability in the absence of titration boxes in their simulations. By contrast, we favor that titration plays a more fundamental role in initiation control, because it is the protein counting device in the protocell and also the Δ4 cells, where DnaA-ATP ↔DnaA-ADP conversion is absent. Furthermore, titration boxes, which are prevalent in bacteria, ensure synchronous initiation (as explained in Section E), and explain as to why bacteria produce significantly more DnaA molecules than necessary for *ori*. Albeit titration is fundamental in our model, its performance is not perfect in terms of initiation instability, and we demonstrated that RIDA is the key conversion element required for initiation stability when titration is in place.

While the details of molecular effects on initiation are beyond the scope of this theory work, we suggest recent work by Elf and colleagues [48] and by us on various deletion mutants, including Δ4 [13], for single-cell level experimental investigation as confirmations of some of our predictions..

### E. Asynchrony and cell-to-cell variability of initiation in the initiator-titration framework

Initiation stability raises a related issue of stochasticity in initiation. In the systems biology literature, “noise” is mainly discussed in the context of stochasticity in gene expression, decomposed into “intrinsic” vs. “extrinsic” components [49–52]. In our view, there are parallel observations in replication initiation: the initiation asynchrony among *ori*’s within the same cell [13, 53, 54], and the cell-to-cell variability of the initiation mass [6, 13, 55], as illustrated in Fig. 6(a). In this section, we discuss their origins and statistical properties within our initiator-titration model v2 framework.

**FIG. 6.**
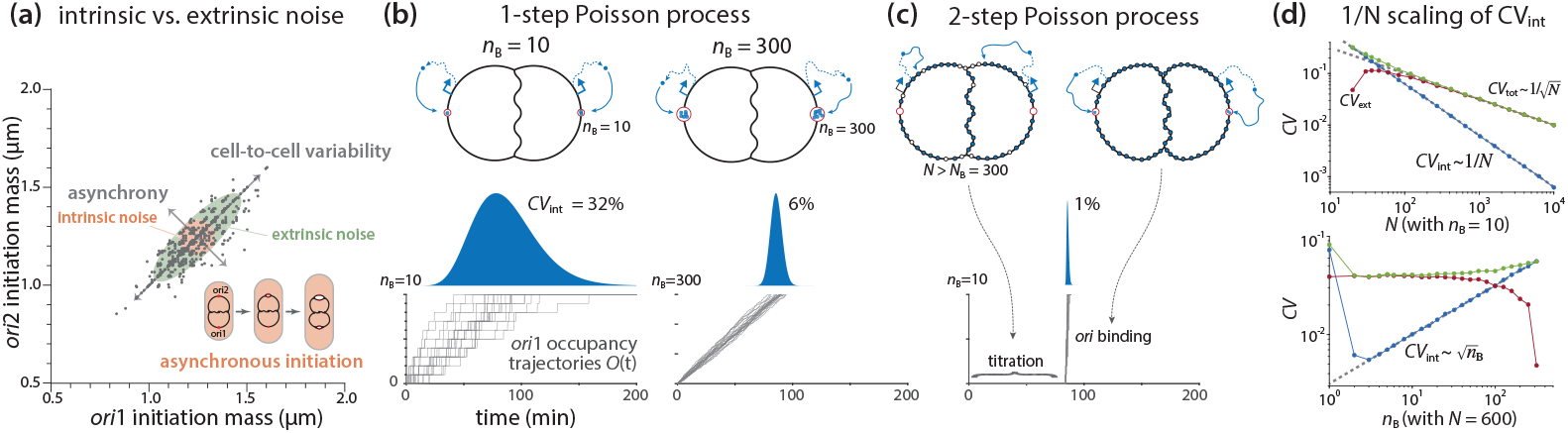
Initiation precision and the reduction in asynchrony by 1*/N* scaling in the two-step Poisson process by titration. (a) Asynchrony (intrinsic noise), extrinsic noise, and cell-to-cell variability in initiation control. Grey dots are single-cell data of wildtype *E. coli* from [13]. (b) Initiation by the first-passage time model based on a simple Poisson process with *n*_B_ the threshold at each *ori*. (left) *n*_B_ = 10 vs. (right) *n*_B_ = 300. (c) Synchronized initiation by titration in the two-step Poisson process. *N* is the mean total number of initiator proteins required for triggering initiation at both *ori*’s. (d) Simulation of the scaling behavior of the intrinsic noise (*CV*_int_), the extrinsic noise (*CV*_ext_), and the total CV (*CV*_tot_) (see Appendix G). Top: varing *N* with fixed *n*_B_; bottom: varing *n*_B_ with fixed *N*. The grey dashed lines are from Eq. 17.

#### 1. Definition of the intrinsic and extrinsic noise

During overlapping cell cycles, the cell contains multiple replication origins at initiation. These origins share the same biochemical environment within one cell, so their initiation events are correlated; the initiation timing in different cells can vary because of stochasticity in biological processes, such as gene expression [49, 52]. On the other hand, since these origins in the same cell do not interact with each other, they can initiate asynchronously due to the innate stochasticity of initiator accumulation at origins [28, 53].

To quantify initiation asynchrony and cell-to-cell variability, we consider two overlapping replication cycles. Suppose the two *ori*’s initiate at initiation mass 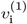 and 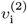, respectively. Similar to stochastic gene expression [49], we can define the intrinsic noise and the extrinsic noise of the initiation mass by the coefficient of variation as

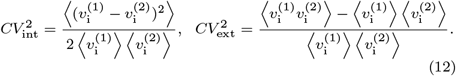

Note that this definition fulfills the relation 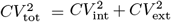, where *CV*_tot_ is the coefficient of variation of the single variable 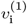 or 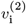 (Appendix F).

We use *CV*_int_ as a measure of asynchrony. Visually, *CV*_int_ describes the width of the off-diagonal axis of the ellipsoid, while *CV*_ext_ describes the elongation extent of the diagonal axis compared to the short axis [Fig. 6(a)]. For example, if *CV*_ext_ = 0, 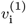 are 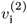 fully uncorrelated, and the ellipsoid becomes a circle. In this case, the intrinsic noise is the sole source of cell-to-cell variability. Generally, while asynchrony is fully determined by the intrinsic noise, the cell-to-cell variability is a result of both the intrinsic noise and the extrinsic noise (see Appendix F for details).

#### 2. A first-passage-time (FPT) model based on a one-step Poisson process

To study the behavior of the extrinsic noise and the intrinsic noise, we convert the initiation mass variables, 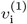 and 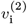, into the first passage time (FPT) variables [56], *T* ^(1)^ and *T* ^(2)^, respectively. That is, the initiator proteins bind to binding sites at *ori*, increasing its occupancy *O*(*t*), and initiate replication as soon as *ori* is fully saturated [*O*(*t*) = *n*_B_]. Although the relation between *v*_i_and FPT is nonlinear, to the zeroth-order approximation, we have

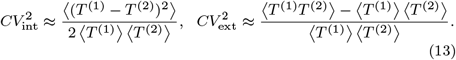

To obtain the scaling law of the noise of FPT, we assume the production of initiator proteins as a Poisson process with a constant production rate *β* [52]. We further assume that all cells are characterized by the same set of physiological parameters without noise. (By this assumption, we are considering the lower bound of the extrinsic noise, and we discuss the contribution of parameter noises in Section E.4.)

Let us first consider a simple scenario of initiation without initiator-titration. In this scenario, there is no chromosomal binding site; all *n*_B_ binding sites are localized at each *ori*, and the initiator protein has an equal probability of binding to either *ori*. That is, the two *ori*’s accumulate the initiator proteins independently. This results in uncorrelated *T* ^(1)^ and *T* ^(2)^ and hence *CV*_ext_ = 0 based on Eq. 13. The intrinsic noise then becomes

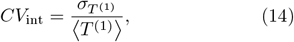

where ⟨*T* ^(1)^⟩ is the mean FPT at *ori* 1 and *σ*_*T*(1)_ is the standard deviation.

In this simplest scenario, the accumulation at *ori*1 is a Poission process with a rate of *β* followed by a binomial trial with equal probability, leading to a Gamma distribution of *T* ^(1)^, with the mean ⟨*T* ^(1)^⟩ = 2*n*_B_*/β* and the standard deviation 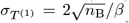 (see Appendix G). Thus, the *CV*_int_ is independent of *β* [56, 57],

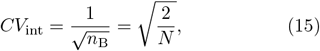

where *N* = 2*n*_B_ is the mean total number of initiator proteins needed for triggering initiation at both *ori*’s (Appendix G).

Therefore, in this one-step Poisson process, the intrinsic noise of FPT scales with the square root of the required total number of initiators *N*. If the number of binding sites at *ori* is *n*_B_ ≈ 10, we have *CV*_int_ ≈ 30% [Fig. 6(b)]. If the cell localizes all *N*_*B*_ ≈ 300 DnaA boxes at each *ori* to increase the threshold, the noise will decrease to 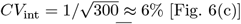 [Fig. 6(c)].

The reason for the 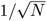 intrinsic noise scaling is that the stochasticity in gene expression fully propagates to the initiation timing, and *T* ^(1)^ and *T* ^(2)^ are uncorrelated. As we explain below, *E. coli* suppresses the intrinsic noise using an ingenious two-step Poisson process by compressing *T* ^(1)^ and *T* ^(2)^ into a narrow range during the cell cycle using titration. In other words, titration of the initiator proteins redirects most gene expression noise to the extrinsic noise, effectively synchronizing *T* ^(1)^ and *T* ^(2)^.

#### 3. A two-step Poisson process in the initiator-titration framework predicts the 1/N scaling of the intrinsic noise, leading to initiation synchrony

Due to the significant differences in the binding affinity between the chromosomal binding sites (*K*_D_ ≈1nM) and *ori* (*K*_D_ ≈100nM), *E. coli* titrates DnaA sequentially in two steps: (1) saturation of the ∼ *N*_B_ chromosomal DnaA boxes by DnaA-ATP and DnaA-ADP, followed by (2) accumulation of DnaA-ATP at *ori* with *n*_B_≪*N*_B_ binding sites. Thus, we modify the one-step Poisson process by adding the titration step, namely, a two-step Poisson process [Fig. 6(c)]. The first step delays the accumulation processes at *ori* 1 and *ori* 2 and they synchronize their initiations, and the intrinsic noise (asynchrony) is a result of stochasticity in the second step.

To analyze the two-step Poisson process, we rewrite the two FPT variables *T* ^(1)^ and *T* ^(2)^ as, *T* ^(1)^ = *T* ^(0)^ +Δ*T* ^(1)^, *T* ^(2)^ = *T* ^(0)^ + Δ*T* ^(2)^. Here, *T* ^(0)^ is the time required to saturate the chromosomal binding sites, whereas Δ*T* ^(1)^ and Δ*T* ^(2)^ denote the additional respective times for the two *ori*’s to accumulate the initiator proteins to trigger initiation. We assume that *T* ^(0)^, Δ*T* ^(1)^, and Δ*T* ^(2)^ are three independent stochastic variables. Specifically, *T* ^(0)^ follows the original Poisson process with an accumulation rate of *β*, while Δ*T* ^(1)^ and Δ*T* ^(2)^ each independently follows the same Poisson process with an accumulation rate of *β/*2 (initiator proteins produced at the rate *β* bind the two *ori*’s), as derived in Appendix G. By this decomposition, Eq. 13 can be rewritten as

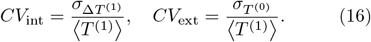

According to the corresponding Gamma distributions, the mean FPT reads ⟨*T* ^(1)^⟩ = *N/β*, where *N* is the mean total number of initiator proteins needed for triggering initiation at both *ori*’s; the standard deviation of the first-step FPT reads 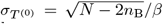, and the standard deviation of the second-step FPT reads 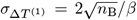 (see Appendix G). Therefore, basedon Eq. 16, we obtain the CV’s scaling law as

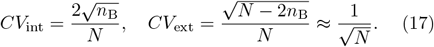

This result indicates that *CV*_int_ decays in ∼ 1*/N*, much faster than the total noise 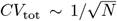, and *CV*_ext_becomes the dominant noise component when *N* is large. For example, if *n*_B_ = 10, *N*_B_ ≈ 300, and *N* ≈ 2(*N*_B_ + *n*_B_) ≈600 (two-overlapping cell cycle), the noise of the two-step processes decreases dramatically from ∼ 30% to only 1%, while the extrinsic noise is around 4%.

To test the predictions of the two-step Poisson process, we conducted simulation by considering a Poissonian protein production followed by a partitioning among three destinations: chromosomal binding sites, *ori* 1, and *ori* 2 (see Appendix G for model settings). As shown in Fig. 6(d), the scaling behavior of the intrinsic noise and the extrinsic noise is consistent with Eq. 17.

In summary, the chromosomal titration boxes effectively synchronize the accumulation of DnaA-ATP at multiple *ori*’s by titration, compressing their initiation timing into a narrow temporal window during the cell cycle [53, 58]. This is consistent with long-standing experimental observations of synchronous initiation of minichromosomes [59, 60], and more recent observations of ectopic chromosomal origins [61]. This improvement of precision by two sequential binding processes is reminiscent of the ratchet-like kinetic proofreading model, and our results are generalizable.

#### 4. Other noise sources not quantified in this work

In the previous section, we have mainly discussed asynchrony and cell-to-cell variability in initiation resulting from stochastic protein production, which predicts *CV*_int_≈ 1%, and *CV*_ext_ ≈ 4% in *E. coli*. However, experimentally measured *CV*_int_ is about 3% -4% [13] and *CV*_tot_ is about 10% [6, 13, 55], both larger than the prediction. For mutants lacking DnaA-ATP↔ DnaA-ADP conversion elements, the cell-to-cell variability can increase up to 20% [13]. The likely sources of additional asynchrony and cell-to-cell variability are likely as follows.

For the intrinsic noise, the initiator accumulation at the two *ori*’s can be negatively correlated because of the new round of replication. Once the first initiation event is triggered at one *ori*, the newly produced DnaA boxes will titrate DnaA, and the newly activated RIDA decreases the DnaA-ATP pool [Fig. 5(b)], further delaying the initiation of the second *ori*. This anti-correlation between two asynchronous initiations should increase the intrinsic noise *CV*_int_.

For the extrinsic noise, we suggest two extra main sources other than the 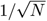 for titration by Poisson process (Eq. 17): (1) cell-to-cell variability in the initiator concentration [52, 62, 63], DnaA-ATP/ADP ratio, doubling time [9, 64], and C period [36, 65], and (2) the growth-condition (*C/τ*) dependent initiation instability discussed in Section D [Figs. 4(c) & 5(c)]. In principle, even without noises in C period and doubling time, instability can cause a bimodal distribution of the initiation mass that significantly increases the extrinsic noise [11]. The noise caused by initiation instability should be significant in mutants without the RIDA mechanism, such as the Δ4 cells [13].

For the quantification of these noise contributions, we leave a more detailed analysis to future work.

## III. CONCLUSION AND PERSPECTIVE

In this work, we have provided a comprehensive quantitative explanation of how bacteria control the cell cycle under balanced growth, particularly focusing on replication initiation as a tractable problem. Our analysis builds upon the original initiator-titration model proposed by Hansen and colleagues [19], which offered valuable insights into the two-step initiation processes that trigger initiation.

Over the past three decades, significant progress has been made in understanding the conserved master replication initiator protein, DnaA. One perplexing aspect has been the coexistence of two forms of DnaA (DnaA-ATP and DnaA-ADP), with only DnaA-ATP being initiation competent. Expanding upon the original model by Hansen and colleagues, we developed the initiator-titration model v2, which incorporates the two-state DnaA model and accounts for DnaA box distribution. We have derived an analytical expression for the initiation mass in terms of three mechanistic parameters for DnaA: its concentration, the average ratio [DnaA-ATP]/[DnaA-ADP], and the number of DnaA titration boxes (Eq. 11). However, through our dynamical stability analysis, we have also revealed a previously unexplored instability in initiation within this model [Fig. 4(c)], thereby elucidating recent observations from numerical simulations by Berger and ten Wolde [11]. We have demonstrated that the replication-dependent DnaA-ATP→DnaA-ADP conversion (by RIDA) alone restores initiation stability [66]. Additionally, when considering all extrinsic DnaA-ATP ↔ DnaA-ADP elements, the initiation mass remains remarkably invariant across a wide range of growth conditions, in agreement with experimental observations [33, 36, 67].

Moreover, we have discovered that the titration process of the chromosomal DnaA boxes suppresses the intrinsic noise or asynchrony in initiation by *CV* ∼ 1*/N* scaling. This finding represents a significant improvement over the naively expected standard coefficient of variation scaling 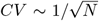 for a Poisson process. It underscores the extraordinary consequences of the twostep initiation processes in the initiator-titration models, highlighting the remarkable precision achieved by bacteria.

In conclusion, we propose that titration may have been a pivotal evolutionary milestone, acting as a protein-counting mechanism that co-evolved with balanced biosynthesis. This system would not only enable bacteria to homeostatically control their size via the adder principle, but also lead to synchronous initiation by effectively separating titration and initiation in two steps. Our results thus illuminate how bacteria employ a seemingly straightforward yet efficient titration-based strategy to address fundamental biological challenges. This differentiates them from eukaryotes, which use programmed gene expression and protein degradation to sense and control protein concentrations. While our findings focus on a specific case of initiation control, they also trigger intriguing questions about the potential pervasiveness of titration-based precision control in diverse biological systems. Uncovering additional examples of such mechanisms will significantly advance our overall understanding of precision control and pave the way for practical applications, including the design of synthetic cells.

## ACKNOWLEDGMENTS

We thank Flemming Hansen, Tove Atlung, Tsutomu Katayama, Anders Lobner-Olesen, Godefroid Charbon, Thias Boesen, Johan Elf, Dongyang Li, Fangwei Si, Guillaume Le Treut, Cara Jensen, Mareike Berger, Pieter-Rein ten Wolde, Alan Leonard, Julia Grimwade, Conrad Woldringh, Charles Helmstetter, and Willie Donachie for many invaluable discussions and exchange of ideas over the years that inspired and helped shape the ideas presented in this work. This work was supported by NSF MCB-2016090 and NIH MIRA (R35GM139622) to SJ.

## Reference

[1] L. Wolpert, Positional information and the spatial pattern of cellular differentiation, J. Theor. Biol. 25, 1 (1969).

[2] T. Evans, E. T. Rosenthal, J. Youngblom, D. Distel, and T. Hunt, Cyclin: a protein specified by maternal mRNA in sea urchin eggs that is destroyed at each cleavage division, Cell 33, 389 (1983).

[3] S. Jun, F. Si, R. Pugatch, and M. Scott, Fundamental principles in bacterial physiology—history, recent progress, and the future with focus on cell size control: a review, Rep. Prog. Phys. 81, 056601 (2018).

[4] M. Schaechter, O. Maaløe, and N. O. Kjeldgaard, Dependency on medium and temperature of cell size and chemical composition during balanced growth of salmonella typhimurium, Microbiology 19, 592 (1958).

[5] N. O. Kjeldgaard, O. Maaloe, and M. Schaechter, The transition between different physiological states during balanced growth of salmonella typhimurium, J. Gen. Microbiol. 19, 607 (1958).

[6] F. Si, G. Le Treut, J. T. Sauls, S. Vadia, P. A. Levin, and S. Jun, Mechanistic origin of Cell-Size control and homeostasis in bacteria, Curr. Biol. 29, 1760 (2019).

[7] T. den Blaauwen, L. W. Hamoen, and P. A. Levin, The divisome at 25: the road ahead, Curr. Opin. Microbiol. (2017).

[8] M. Campos, I. V. Surovtsev, S. Kato, A. Paintdakhi, B. Beltran, S. E. Ebmeier, and C. Jacobs-Wagner, A constant size extension drives bacterial cell size homeostasis, Cell 159, 1433 (2014).

[9] S. Taheri-Araghi, S. Bradde, J. T. Sauls, N. S. Hill, P. A. Levin, J. Paulsson, M. Vergassola, and S. Jun, Cell-Size control and homeostasis in bacteria, Curr. Biol. 25, 385 (2015).

[10] S. Jun and S. Taheri-Araghi, Cell-size maintenance: universal strategy revealed, Trends Microbiol. 23, 4 (2015).

[11] M. Berger and P. R. T. Wolde, Robust replication initiation from coupled homeostatic mechanisms, Nat. Commun. 13, 6556 (2022).

[12] L. Sompayrac and O. Maaloe, Autorepressor model for control of DNA replication, Nat. New Biol. 241, 133 (1973).

[13] T. Boesen, G. Charbon, H. Fu, C. Jensen, D. Li, S. Jun, and others, Robust control of replication initiation in the absence of DnaA-ATP DnaA-ADP regulatory elements in escherichia coli, bioRxiv (2022).

[14] A. C. Leonard, P. Rao, R. P. Kadam, and J. E. Grimwade, Changing perspectives on the role of DnaA-ATP in orisome function and timing regulation, Front. Microbiol. 10, 2009 (2019).

[15] F. G. Hansen and T. Atlung, The DnaA tale, Front. Microbiol. 9, 319 (2018).

[16] T. Katayama, K. Kasho, and H. Kawakami, The DnaA cycle in escherichia coli: Activation, function and inactivation of the initiator protein, Front. Microbiol. 8, 2496 (2017).

[17] L. Riber, J. Frimodt-Møller, G. Charbon, and A. Løbner-Olesen, Multiple DNA binding proteins contribute to timing of chromosome replication in e. coli, Front Mol Biosci 3, 29 (2016).

[18] T. Katayama, S. Ozaki, K. Keyamura, and K. Fujimitsu, Regulation of the replication cycle: conserved and diverse regulatory systems for DnaA and oric, Nat. Rev. Microbiol. 8, 163 (2010).

[19] F. G. Hansen, B. B. Christensen, and T. Atlung, The initiator titration model: computer simulation of chromosome and minichromosome control, Res. Microbiol. 142, 161 (1991).

[20] T. Katayama, T. Kubota, K. Kurokawa, E. Crooke, and K. Sekimizu, The initiator function of DnaA protein is negatively regulated by the sliding clamp of the e. coli chromosomal replicase, Cell 94, 61 (1998).

[21] J. Kato and T. Katayama Hda, a novel DnaA-related protein, regulates the replication cycle in escherichia coli, EMBO J. 20, 4253 (2001).

[22] A. C. Leonard and J. E. Grimwade, The orisome: structure and function, Front. Microbiol. 6, 545 (2015).

[23] K. Sekimizu, D. Bramhill, and A. Kornberg, ATP activates dnaa protein in initiating replication of plasmids bearing the origin of the e. coli chromosome, Cell 50, 259 (1987).

[24] S. Nishida, K. Fujimitsu, K. Sekimizu, T. Ohmura, T. Ueda, and T. Katayama, A nucleotide switch in the escherichia coli DnaA protein initiates chromosomal replication: evidence from a mutant DnaA protein defective in regulatory ATP hydrolysis in vitro and in vivo, J. Biol. Chem. 277, 14986 (2002).

[25] K. C. McGarry, V. T. Ryan, J. E. Grimwade, and A. C. Leonard, Two discriminatory binding sites in the escherichia coli replication origin are required for DNA strand opening by initiator DnaA-ATP, Proc. Natl. Acad. Sci. U. S. A. 101, 2811 (2004).

[26] S. Schaper and W. Messer, Interaction of the initiator protein DnaA of escherichia coli with its DNA target, J. Biol. Chem. 270, 17622 (1995).

[27] H. Kawakami, K. Keyamura, and T. Katayama, Formation of an ATP-DnaA-specific initiation complex requires DnaA arginine 285, a conserved motif in the AAA+ protein family, J. Biol. Chem. 280, 27420 (2005).

[28] J. Herrick, M. Kohiyama, T. Atlung, and F. G. Hansen, The initiation mess?, Mol. Microbiol. 19, 659 (1996).

[29] M. H. Buckstein, J. He, and H. Rubin, Characterization of nucleotide pools as a function of physiological state in escherichia coli, J. Bacteriol. 190, 718 (2008).

[30] L. S. Hsieh, J. Rouviere-Yaniv, and K. Drlica, Bacterial DNA supercoiling and [ATP]/[ADP] ratio: changes associated with salt shock, J. Bacteriol. 173, 3914 (1991).

[31] More precisely, there are 11 DnaA boxes are clustered within ori, with three high-affinity sites and eight low-affinity sites. Only DnaA-ATP can bind to the eight low-affinity sites. For simplicity, we used “10” to denote the number of low-affinity binding sites at ori throughout this paper.

[32] F. G. Hansen, T. Atlung, R. E. Braun, A. Wright, P. Hughes, and M. Kohiyama, Initiator (DnaA) protein concentration as a function of growth rate in escherichia coli and salmonella typhimurium, J. Bacteriol. 173, 5194 (1991).

[33] H. Zheng, Y. Bai, M. Jiang, T. A. Tokuyasu, X. Huang, F. Zhong, Y. Wu, X. Fu, N. Kleckner, T. Hwa, and C. Liu, General quantitative relations linking cell growth and the cell cycle in escherichia coli, Nat Microbiol 5, 995 (2020).

[34] T. M. Pham, K. W. Tan, Y. Sakumura, K. Okumura, H. Maki, and M. T. Akiyama, A single-molecule approach to DNA replication in escherichia coli cells demonstrated that DNA polymerase III is a major determinant of fork speed, Mol. Microbiol. 90, 584 (2013).

[35] D. Bhat, S. Hauf, C. Plessy, Y. Yokobayashi, and S. Pigolotti, enCorrection: Speed variations of bacterial replisomes, Elife 12 (2023).

[36] F. Si, D. Li, S. E. Cox, J. T. Sauls, O. Azizi, C. Sou, A. B. Schwartz, M. J. Erickstad, Y. Jun, X. Li, and S. Jun, Invariance of initiation mass and predictability of cell size in escherichia coli, Curr. Biol. 27, 1278 (2017).

[37] S. Cooper and C. E. Helmstetter, Chromosome replication and the division cycle of escherichia colibr, J. Mol. Biol. (1968).

[38] R. Kitagawa, H. Mitsuki, T. Okazaki, and T. Ogawa, A novel DnaA protein-binding site at 94.7 min on the escherichia coli chromosome, Mol. Microbiol. 19, 1137 (1996).

[39] K. Kasho and T. Katayama, DnaA binding locus datA promotes DnaA-ATP hydrolysis to enable cell cycle-coordinated replication initiation, Proceedings of the National Academy of Sciences 110, 936 (2013).

[40] K. Fujimitsu, T. Senriuchi, and T. Katayama, Specific genomic sequences of e. coli promote replicational initiation by directly reactivating ADP-DnaA, Genes Dev. 23, 1221 (2009).

[41] K. Kasho, K. Fujimitsu, T. Matoba, T. Oshima, and T. Katayama, Timely binding of IHF and fis to DARS2 regulates ATP-DnaA production and replication initiation, Nucleic Acids Res. 42, 13134 (2014).

[42] M. C. Moolman, S. T. Krishnan, J. W. J. Kerssemakers, A. van den Berg, P. Tulinski, M. Depken, R. Reyes-Lamothe, D. J. Sherratt, and N. H. Dekker, Slow unloading leads to DNA-bound β2-sliding clamp accumulation in live escherichia coli cells, Nat. Commun. 5, 5820 (2014).

[43] M. Lu, J. L. Campbell, E. Boye, and N. Kleckner, SeqA: a negative modulator of replication initiation in e. coli, Cell 77, 413 (1994).

[44] F. G. Hansen, B. B. Christensen, C. B. Nielsen, and T. Atlung, Insights into the quality of DnaA boxes and their cooperativity, J. Mol. Biol. 355, 85 (2006).

[45] T. A. Rozgaja, J. E. Grimwade, M. Iqbal, C. Czerwonka, M. Vora, and A. C. Leonard, Two oppositely oriented arrays of low-affinity recognition sites in oric guide progressive binding of DnaA during escherichia coli pre-RC assembly, Mol. Microbiol. 82, 475 (2011).

[46] W. D. Donachie and G. W. Blakely, Coupling the initiation of chromosome replication to cell size in escherichia coli, Curr. Opin. Microbiol. 6, 146 (2003).

[47] K. Kurokawa, S. Nishida, A. Emoto, K. Sekimizu, and T. Katayama, Replication cycle-coordinated change of the adenine nucleotide-bound forms of DnaA protein in escherichia coli, EMBO J. 18, 6642 (1999).

[48] A. Knöppel, O. Broström, K. Gras, J. Elf, and D. Fange, Regulatory elements coordinating initiation of chromosome replication to the Escherichia coli cell cycle, Proceedings of the National Academy of Sciences 120, e2213795120 (2023).

[49] M. B. Elowitz, A. J. Levine, E. D. Siggia, and P. S. Swain, Stochastic gene expression in a single cell, Science 297, 1183 (2002).

[50] M. Thattai and A. van Oudenaarden, Intrinsic noise in gene regulatory networks, Proc. Natl. Acad. Sci. U. S. A. 98, 8614 (2001).

[51] J. Paulsson, Summing up the noise in gene networks, Nature 427, 415 (2004).

[52] Y. Taniguchi, P. J. Choi, G.-W. Li, H. Chen, M. Babu, J. Hearn, A. Emili, and X. Sunney Xie, Quantifying e. coli proteome and transcriptome with Single-Molecule sensitivity in single cells,Science (2010).

[53] K. Skarstad, E. Boye, and H. B. Steen, Timing of initiation of chromosome replication in individual escherichia coli cells, EMBO J. 5, 1711 (1986).

[54] A. Løbner-Olesen, K. Skarstad, F. G. Hansen, K. von Meyenburg, and E. Boye, The DnaA protein determines the initiation mass of escherichia coli K-12, Cell 57, 881 (1989).

[55] J. T. Sauls, S. E. Cox, Q. Do, V. Castillo, Z. Ghulam-Jelani, and S. Jun, Control of bacillus subtilis replication initiation during physiological transitions and perturbations, MBio 10 (2019).

[56] P. P. Pandey, H. Singh, and S. Jain, Exponential trajectories, cell size fluctuations, and the adder property in bacteria follow from simple chemical dynamics and division control, Phys. Rev. E 101, 062406 (2020).

[57] K. R. Ghusinga, J. J. Dennehy, and A. Singh, First-passage time approach to controlling noise in the timing of intracellular events, Proceedings of the National Academy of Sciences 114, 693 (2017).

[58] M. Wallden, D. Fange, E. G. Lundius, Ö. Baltekin, and J. Elf, The synchronization of replication and division cycles in individual e. coli cells, Cell 166, 729 (2016).

[59] A. C. Leonard and C. E. Helmstetter, Cell cycle-specific replication of escherichia coli minichromosomes, Proc. Natl. Acad. Sci. U. S. A. 83, 5101 (1986).

[60] C. E. Helmstetter and A. C. Leonard, Coordinate initiation of chromosome and minichromosome replication in escherichia coli, J. Bacteriol. 169, 3489 (1987).

[61] X. Wang, C. Lesterlin, R. Reyes-Lamothe, G. Ball, and D. J. Sherratt, Replication and segregation of an escherichia coli chromosome with two replication origins, Proc. Natl. Acad. Sci. U. S. A. 108, E243 (2011).

[62] H. Salman, N. Brenner, C.-K. Tung, N. Elyahu,E. Stolovicki, L. Moore, A. Libchaber, and E. Braun, Universal protein fluctuations in populations of microorganisms, Phys. Rev. Lett. 108, 238105 (2012).

[63] P. Wang, L. Robert, J. Pelletier, W. L. Dang, F. Taddei Wright, and S. Jun, Robust growth of escherichia coli, Curr. Biol. 20, 1099 (2010).

[64] M. Schaechter, J. P. Williamson, J. R. Hood, Jr, and A. L. Koch, Growth, cell and nuclear divisions in some bacteria, J. Gen. Microbiol. 29, 421 (1962).

[65] G. Le Treut, F. Si, D. Li, and S. Jun, Quantitative examination of five stochastic Cell-Cycle and Cell-Size control models for escherichia coli and bacillus subtilis, Front. Microbiol. 12, 721899 (2021).

[66] L. Riber, J. A. Olsson, R. B. Jensen, O. Skovgaard, S. Dasgupta, M. G. Marinus, and A. Løbner-Olesen, Hda-mediated inactivation of the DnaA protein and dnaa gene autoregulation act in concert to ensure homeostatic maintenance of the escherichia coli chromosome, Genes Dev. 20, 2121 (2006).

[67] W. D. Donachie, Relationship between cell size and time of initiation of DNA replication, Nature 219, 1077 (1968).

