## Appendix for "Replication initiation in bacteria: precision control based on protein counting"

Haochen Fu and Fangzhou Xiao
*Department of Physics, University of California San Diego, 9500 Gilman Dr, La Jolla, CA 92093*

Suckjoon Jun
*Department of Physics and Department of Molecular Biology,*
*University of California San Diego, 9500 Gilman Dr, La Jolla, CA 92093*

#### Appendix A: Derivation of the steady-state initiation mass formula in the protocell model

We consider one generation from cell birth to cell division in the steady state. First, due to exponential growth and balanced biosynthesis of DnaA, we have

$$V(t) = V(0)e^{\lambda t}, \tag{A1}$$

$$I(t) = I(0)e^{\lambda t}, \tag{A2}$$

where $V(t)$ is the cell volume, $I(t)$ is the number of initiators, $\lambda = \frac{\ln 2}{\tau}$ is the growth rate, and $0 \leq t \leq \tau$.

We denote the number of chromosomal binding sites as $\tilde{B}(t)$. The behavior of $\tilde{B}(t)$ is complicated and depends on the ratio of $C/\tau$. Suppose $n\tau \leq C \leq (n+1)\tau$, where $n \equiv \lfloor \frac{C}{\tau} \rfloor$. The shape of $\tilde{B}(t)$ also depends on the relative timing of initiation and termination. For example, Fig. 1A shows when $n = 0$ and $t_{\text{ini}} > t_{\text{ter}}$, and Fig. 1B shows when $n = 1$ and $t_{\text{ini}} < t_{\text{ter}}$. We will discuss the two cases $t_{\text{ini}} \leq t_{\text{ter}}$ and $t_{\text{ini}} > t_{\text{ter}}$ separately.

##### 1. $t_{\text{ini}} \leq t_{\text{ter}}$

As illustrated in Fig. 1C, in this case, the termination time is determined by $\tau$, $C$, and $t_{\text{ini}}$ as

$$t_{\text{ter}} = t_{\text{ini}} + C - n\tau. \tag{A3}$$

The condition for this situation is $t_{\text{ter}} \leq \tau$, which gives

$$t_{\text{ini}} \leq (n+1)\tau - C. \tag{A4}$$

As shown in Fig. 1C, the curve $\tilde{B}(t)$ consists of three segments with different slopes. By mathematical induction, we can obtain the expression of each slope,

$$k_1 = (2^n - 1)k, \qquad 0 \leq t < t_{\text{ini}}, \tag{A5a}$$

$$k_2 = (2^{n+1} - 1)k, \quad t_{\text{ini}} \leq t < t_{\text{ter}}, \tag{A5b}$$

$$k_3 = (2^{n+1} - 2)k, \quad t_{\text{ter}} \leq t \leq \tau, \tag{A5c}$$

where $k$ is the replication rate of the DnaA boxes by a pair of replication forks, i.e., following to Assumptions 3 & 4,

$$k \equiv \frac{N_{\text{B}}}{C}. \tag{A6}$$

The initial value $\tilde{B}(0)$ is $\tilde{B}(\tau)/2$, hence we have

$$\tilde{B}(0) = k_1 t_{\text{ini}} + k_2(t_{\text{ter}} - t_{\text{ini}}) + k_3(\tau - t_{\text{ter}}).$$

Using Eqs. A3 and A5, we obtain

$$\tilde{B}(0) = N_{\text{B}} \left[ 1 + (2^{n+1} - n - 2)\frac{\tau}{C} - (2^n - 1)\frac{t_{\text{ini}}}{C} \right]. \tag{A7}$$

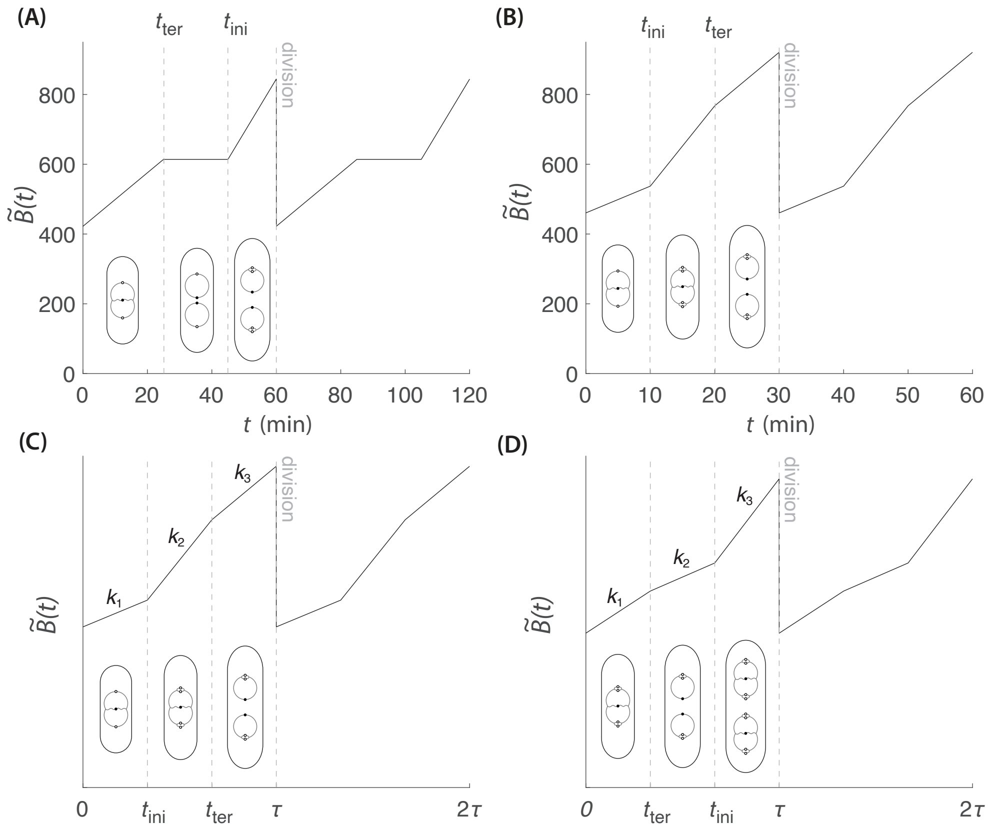


FIG. 1. The time trajectory of the number of chromosomal initiator binding sites (titration boxes) $\tilde{B}(t)$ in different scenarios in the steady state. (a) $n = 0$ and $t_{\rm ini} > t_{\rm ter}$. (b) $n = 1$ and $t_{\rm ini} < t_{\rm ter}$. (c) The general case when $t_{\rm ini} \leq t_{\rm ter}$. (d) The general case when $t_{\rm ini} > t_{\rm ter}$.

Once we obtain the expressions of $\tilde{B}(0)$, the slopes $k_1$, $k_2$, and $k_3$, $\tilde{B}(t)$ is fully determined. Of our particular interest, we have

$$\tilde{B}(t_{\rm ini}) = N_{\rm B}\left[1 + (2^{n+1} - n - 2)\frac{\tau}{C}\right], \tag{A8}$$

which is independent of $t_{\rm ini}$.

#### 2. $t_{\rm ini} > t_{\rm ter}$

As illustrated in Fig. 1D, in this case, the termination time can be written as

$$t_{\rm ter} = t_{\rm ini} + C - (n+1)\tau. \tag{A9}$$

The condition for this situation is $t_{\rm ter} \geq 0$, which gives

$$t_{\rm ini} > (n+1)\tau - C. \tag{A10}$$

Similarly, we obtain

$$k_1 = (2^{n+1} - 1)k, \quad 0 \leq t < t_{\rm ter}, \tag{A11a}$$

$$k_2 = (2^{n+1} - 2)k, \quad t_{\rm ter} \leq t < t_{\rm ini}, \tag{A11b}$$

$$k_3 = (2^{n+2} - 2)k, \quad t_{\rm ini} \leq t \leq \tau, \tag{A11c}$$

and the initial value $\tilde{B}(0)$ reads

$$\begin{aligned}\tilde{B}(0) &= k_1 t_{\text{ter}} + k_2(t_{\text{ini}} - t_{\text{ter}}) + k_3(\tau - t_{\text{ini}}) \\ &= N_{\text{B}}\left[1 + (2^{n+2} - n - 3)\frac{\tau}{C} - (2^{n+1} - 1)\frac{t_{\text{ini}}}{C}\right].\end{aligned} \tag{A12}$$

Likewise, Eqs. A11 and A12 allow us to calculate $\tilde{B}(t)$ at any time. In particular, at initiation,

$$\tilde{B}(t_{\text{ini}}) = 2N_{\text{B}}\left[1 + (2^{n+1} - n - 2)\frac{\tau}{C}\right]. \tag{A13}$$

which is again independent of $t_{\text{ini}}$.

#### 3. Initiation mass formula

Now $\tilde{B}(t)$ can be fully determined by the given parameters. Let us go back to Eqs. A1 and A2. To calculate the initiation mass, we only need to calculate $I(t_{\text{ini}})$. Based on our initiation criteria, $I(t_{\text{ini}}) = B(t_{\text{ini}}) = \tilde{B}(t_{\text{ini}}) + n_{\text{B}} \cdot \#ori$, we have

$$I(t_{\text{ini}}) = \begin{cases} \tilde{B}(t_{\text{ini}}) + 2^n n_{\text{B}}, & t_{\text{ini}} \leq (n+1)\tau - C, \\ \tilde{B}(t_{\text{ini}}) + 2^{n+1} n_{\text{B}}, & t_{\text{ini}} > (n+1)\tau - C, \end{cases} \tag{A14}$$

The cell volume at initiation reads

$$V(t_{\text{ini}}) = \frac{I(t_{\text{ini}})}{c_{\text{I}}}, \tag{A15}$$

and the initiation mass, by definition, reads

$$v_{\text{i}} \equiv \frac{V(t_{\text{ini}})}{\#ori} = \begin{cases} \frac{V(t_{\text{ini}})}{2^n}, & t_{\text{ini}} \leq (n+1)\tau - C, \\ \frac{V(t_{\text{ini}})}{2^{n+1}}, & t_{\text{ini}} > (n+1)\tau - C. \end{cases} \tag{A16}$$

By plugging Eqs. A15, A14, A8 and A13 into this formula, we obtain the full expression of the initiation mass

$$v_{\text{i}} = \frac{1}{c_{\text{I}}}\left\{\left[\frac{1}{2^n} + (2 - \frac{n+2}{2^n})\frac{\tau}{C}\right] N_{\text{B}} + n_{\text{B}}\right\}, \quad n = \lfloor\frac{C}{\tau}\rfloor. \tag{A17}$$

This is exactly Eq. 2 in Section B in the main text. As we can see, the initiation mass is independent of the initiation time $t_{\text{ini}}$.

### Appendix B: Derivation of the stability regimes for the initiation mapping $\mathcal{F}$

We start from the initiation mapping $\mathcal{F} : \mathbb{R}^{d-1} \to \mathbb{R}^d, \boldsymbol{\rho} \mapsto \boldsymbol{\rho}^+$ derived in Section C in the main text,

$$\rho_i^+ = \begin{cases} \rho_{i-1} + \frac{t^+}{C}, & \text{if} \quad \rho_{i-1} + \frac{t^+}{C} < 1, \\ 1, & \text{else}, \end{cases} \tag{B1}$$

where the initiation time $t^+$ is determined by

$$N_{\text{B}}\left(2^{-d} + \sum_{i=1}^{d} \rho_i^+ 2^{-i}\right) + n_B = \frac{e^{\lambda t^+}}{2}\left[N_{\text{B}}\left(2^{-(d-1)} + \sum_{i=1}^{d-1} \rho_i 2^{-i}\right) + n_B\right]. \tag{B2}$$

#### 1. Steady state

First, note that for any finite $d$, the image space of the map $\mathcal{F}$ is different from its preimage space, so there is no fixed point of $\mathcal{F}$. However, we can consider the $d \to \infty$ limit. The reason is as follows.

Consider the left-hand side of Eq. B2. Due to the definition of $\rho_i$, if $d \to \infty$, there exists $n \geq 0$, s.t., $\rho_{n+1} = 1$, and we have

$$\begin{aligned} N_{\mathrm{B}}\left(2^{-d}+\sum_{i=1}^{d}\rho_i 2^{-i}\right) &= N_{\mathrm{B}}\left(2^{-d}+\sum_{i=1}^{n}\rho_i 2^{-i}+\sum_{i=n+1}^{d}2^{-i}\right) \\ &= N_{\mathrm{B}}\left(2^{-d}+\sum_{i=1}^{n}\rho_i 2^{-i}+2^{-n}-2^{-d}\right) \\ &= N_{\mathrm{B}}\left(2^{-n}+\sum_{i=1}^{n}\rho_i 2^{-i}\right). \end{aligned} \tag{B3}$$

This indicates that if $d$ is large enough, we can seek the "fixed-point"-like solution that

$$\rho_i^{+} = \rho_i, \quad \text{if} \quad \rho_i < 1. \tag{B4}$$

Plugging this solution into Eq. B2 and taking Eq. B3 into consideration, we obtain

$$e^{\lambda t_{\mathrm{ss}}^{+}} = 2, \quad \text{or} \quad t_{\mathrm{ss}}^{+} = \tau. \tag{B5}$$

This is exactly the steady-state periodic assumption in Appendix A. Thus, the steady-state solution with a period of $\tau$ is mathmatically equivalent to the "fixed-point" solution when $d \to \infty$. According to Eqs. B1 and B4, we obtain

$$\rho_i^{\mathrm{ss}} = \begin{cases} i\frac{\tau}{C}, & \text{if} \quad i \leq n, \\ 1, & \text{else}, \end{cases} \tag{B6}$$

where

$$n \equiv \lfloor \frac{C}{\tau} \rfloor, \tag{B7}$$

is the number of overlapping replication cycles in the steady state. In the steady state, we can also calculate the number of initiators at initiation $I(t_{\mathrm{ini}})$,

$$\begin{aligned} I(t_{\mathrm{ini}}) &= B(t_{\mathrm{ini}}) \\ &= N_{\mathrm{B}}\left(2^{d}+\sum_{i=1}^{d}\rho_i^{\mathrm{ss}}2^{d-i}\right)+2^{d}n_{\mathrm{B}} \\ &= 2^{d}\left[N_{\mathrm{B}}\left(2^{-d}+\sum_{i=1}^{d}\rho_i^{\mathrm{ss}}2^{-i}\right)+n_{\mathrm{B}}\right] \\ &= 2^{d}\left[N_{\mathrm{B}}\left(2^{-n}+\sum_{i=1}^{n}\rho_i^{\mathrm{ss}}2^{-i}\right)+n_{\mathrm{B}}\right] \\ &= 2^{d}\left[N_{\mathrm{B}}\left(2^{-n}+\frac{\tau}{C}\sum_{i=1}^{n}i2^{-i}\right)+n_{\mathrm{B}}\right] \\ &= 2^{d}\left\{N_{\mathrm{B}}\left[2^{-n}+(2-\frac{n+2}{2^{n}})\frac{\tau}{C}\right]+n_{\mathrm{B}}\right\}. \end{aligned} \tag{B8}$$

The steady-state initiation mass $v_{\mathrm{i}}^{\mathrm{ss}}$ is then given by

$$v_{\mathrm{i}}^{\mathrm{ss}} = \frac{I(t_{\mathrm{ini}})}{2^{d}c_{\mathrm{I}}},$$

which is the same as Eq. A17.

#### 2. The Jacobian matrix at the steady state

To analyze the stability of the steady state above, we need to compute the Jacobian matrix at the steady state,

$$J = \left.\frac{\partial \rho_i^+}{\partial \rho_j}\right|_{\mathrm{ss}}. \tag{B9}$$

Stability requires that the largest magnitude of the eigenvalues of the Jacobian matrix is smaller than 1.

Based on Eqs. B1 and B6, at the steady state, $J$ is reduced to an $n$-dimensional matrix since other derivatives are zeros. Thus, we have

$$\left.\frac{\partial \rho_i^+}{\partial \rho_j}\right|_{\mathrm{ss}} = \delta_{i-1,j} + \frac{1}{C}\left.\frac{\partial t^+}{\partial \rho_j}\right|_{\mathrm{ss}}, \quad i,j = 1,2,\cdots n. \tag{B10}$$

where $\delta_{i,j}$ is the Kronecker delta.

The partial derivatives of $t^+$ can be computed by taking the partial derivatives at both sides of Eq. B2, and substituting Eq. B1 and the steady-state values. The result reads

$$\frac{1}{C}\left.\frac{\partial t^+}{\partial \rho_n}\right|_{\mathrm{ss}} = \left[2^n(1-2\ln 2) + (n+2)\ln 2 - 1 - \ln 2\left(1+2^n\frac{n_\mathrm{B}}{N_\mathrm{B}}\right)\frac{C}{\tau}\right]^{-1} \equiv a, \tag{B11a}$$

$$\frac{1}{C}\left.\frac{\partial t^+}{\partial \rho_i}\right|_{\mathrm{ss}} = 2^{n-i-1}a, \quad i = 1,2,\cdots,n-1. \tag{B11b}$$

Hence, the Jacobian matrix has the following form,

$$J = \begin{pmatrix} 2^{n-2}a & 2^{n-3}a & \cdots & 2a & a & a \\ 1+2^{n-2}a & 2^{n-3}a & \cdots & 2a & a & a \\ 2^{n-2}a & 1+2^{n-3}a & \cdots & 2a & a & a \\ \vdots & \vdots & \ddots & \vdots & \vdots & \vdots \\ 2^{n-2}a & 2^{n-3}a & \cdots & 1+2a & a & a \\ 2^{n-2}a & 2^{n-3}a & \cdots & 2a & 1+a & a \end{pmatrix}. \tag{B12}$$

It may look difficult to write down a general characteristic equation for this matrix. In the following sections, we will discuss the situations when $n \leq 3$, which already covers almost all the experimental growth conditions in *E. coli*.

#### 3. Stability regimes

##### *a. $n = 0$ ($C < \tau$), slow and intermediate growth conditions*

In this simplest case, $\rho_i^+ = 1$ near the steady state, which means the map $mathcalF$ is a constant map, i.e., $J = 0$. Thus, the steady state is always stable.

##### *b. $n = 1$ ($\tau \leq C < 2\tau$), fast growth conditions*

In this case,

$$J = a = \left[1 - \ln 2 - \ln 2\left(1+2\frac{n_\mathrm{B}}{N_\mathrm{B}}\right)\frac{C}{\tau}\right]^{-1}.$$

The stability condition is $|a| < 1$, which gives out

$$\frac{n_\mathrm{B}}{N_\mathrm{B}} > \left(\frac{1}{\ln 2} - \frac{1}{2}\right)\frac{\tau}{C} - \frac{1}{2}. \tag{B13}$$

Note that at $\tau \to C$, there is a critical point for $n_\mathrm{B}$,

$$n_\mathrm{B} = n_{\mathrm{B},1} = \left(\frac{1}{\ln 2} - 1\right)N_\mathrm{B} \approx 0.44 N_\mathrm{B}.$$

If $n_\mathrm{B} > n_{\mathrm{B},1}$, the steady state is always stable in this regime.

*c. $n = 2$ ($2\tau \le C < 3\tau$), very fast growth conditions*

In this case,

$$J = \begin{pmatrix} a & a \\ 1+a & a \end{pmatrix},$$

where

$$a = \left[3 - 4\ln 2 - \ln 2\left(1 + 4\frac{n_{\rm B}}{N_{\rm B}}\right)\frac{C}{\tau}\right]^{-1}.$$

Note that since $n_{\rm B}/N_{\rm B} \ge 0$ and $2 \le C/\tau < 3$, we have $a \in (-1, 0)$. The characteristic equation for $J$ reads

$$\lambda^2 - 2a\lambda - a = 0,$$

which has two imaginary roots,

$$\lambda_\pm = a \pm i\sqrt{-a^2 - a}.$$

Stability requires

$$|\lambda_\pm| = -a < 1.$$

Thus, we obtain the condition,

$$\frac{n_{\rm B}}{N_{\rm B}} > \left(\frac{1}{\ln 2} - 1\right)\frac{\tau}{C} - \frac{1}{4}. \tag{B14}$$

However, since $\frac{\tau}{C} \in (\frac{1}{3}, \frac{1}{2}]$, the right hand side of the inequality is always negative, which means the steady state is always stable for any positive $n_{\rm B}$ in this regime.

*d. $n = 3$ ($3\tau \le C < 4\tau$), extremely fast growth conditions*

In this case,

$$J = \begin{pmatrix} 2a & a & a \\ 1+2a & a & a \\ 2a & 1+a & a \end{pmatrix},$$

where

$$a = \left[7 - 11\ln 2 - \ln 2\left(1 + 8\frac{n_{\rm B}}{N_{\rm B}}\right)\frac{C}{\tau}\right]^{-1}.$$

According to the range of $C$ and $n_{\rm B}$, we have $a \in (-3, 0)$. The characteristic equation for $J$ reads

$$\lambda^3 - 4a\lambda^2 - 2a\lambda - a = 0.$$

By computing the discriminant of this cubic equation, we know that it has one real root $\lambda = \lambda_1$ and two imaginary roots $\lambda = \xi \pm i\theta$. The critical situation has three possibilities: $\lambda_1 = 1$, $\lambda_1 = -1$ and $\xi^2 + \theta^2 = 1$. We find that only $\lambda_1 = -1$ satisfies the range of $a$. Thus, the stability boundary requires $a = -1/3$. Further perturbation analysis gives out the inequality of stability condition,

$$a > -\frac{1}{3},$$

leading to

$$\frac{n_{\rm B}}{N_{\rm B}} > \left(\frac{5}{4\ln 2} - \frac{11}{8}\right)\frac{\tau}{C} - \frac{1}{8}. \tag{B15}$$

Note that at $\tau \to C/3$, there is another critical point for $n_{\rm B}$,

$$n_{\rm B} = n_{\rm B,2} = \left(\frac{5}{\ln 2} - 7\right)\frac{N_{\rm B}}{12} \approx 0.018 N_{\rm B}.$$

If $n_{\rm B} > n_{\rm B,2}$, the steady state is always stable in this regime.

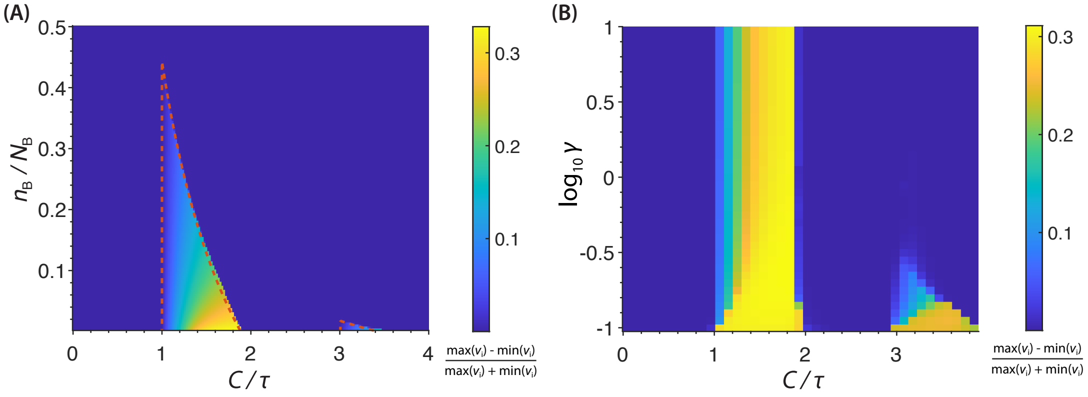


FIG. 2. Simulation of the instability on phase diagrams. Instability is quantified by $(\max v_{\rm i} - \min v_{\rm i})/(\max v_{\rm i} + \min v_{\rm i})$ after 100 doubling times. (a) In the protocell, the unstable regimes match the theoretical predicted unstable regimes (boundaries shown by red dashed curves). (b) In the initiator-titration model v2 with a constant $\gamma$, we fixed $n_B/N_B = 1/30$, and simulate the unstable regimes on the phase diagram of $\gamma$-$\frac{C}{\tau}$.

### Appendix C: A simple derivation for the fixed points of mapping $\mathcal{F}^{\rm o2}$ in $n = 1$

In principle, we can analyze the fixed points of $\mathcal{F}^{\rm o2} \equiv \mathcal{F} \circ \mathcal{F}$ based on the expression of $\mathcal{F}$ (Eqs. B1 and B2). However, since we know from simulation [Fig. 4(c)] that one of the fixed point of $\mathcal{F}^{\rm o2}$ corresponds to

$$v_{\rm i}^{(1)} = \frac{1}{c_{\rm I}}(\alpha^{(1)} N_{\rm B} + n_{\rm B}), \quad \alpha^{(1)} = 1, \tag{C1}$$

we can derive the second fixed point $\alpha^{(2)}$ based on it.

Suppose the initiation event with $\alpha^{(1)} = 1$ at $t = 0$. The initiator number at intiation reads

$$I(0) = 2(N_{\rm B} + n_{\rm B}). \tag{C2}$$

After initiation, $I(t)$ increases exponentially, whereas $B(t)$ first jumps by $2n_{\rm B}$ and then increases linearly with a slope of $2N_B/C$ for a duration time of $C$, as discussed in Section B in the main text. Suppose the next initiation time is at $t = t^+$. According to the initiation criterion $I(t^+) = B(t^+)$, we have

$$2(N_{\rm B} + n_{\rm B})2^{\frac{t^+}{\tau}} = 2(N_{\rm B} + n_{\rm B}) + 2n_{\rm B} + \frac{2N_{\rm B}}{C} t^+, \tag{C3}$$

which provides a transcendental equation to determine $t^+$,

$$2^{\frac{t^+}{\tau}} = 2 + \frac{N_{\rm B}}{N_{\rm B} + n_{\rm B}} \left( \frac{t^+}{C} - 1 \right). \tag{C4}$$

By drawing the curves for the left-hand side and the right-hand side, it is easy to see that $t^+ < \tau$, and $t^+$ exists only when $C > \tau$, which is consistent with the instability condition. Thus, based on Eq. C3, the initiation mass at the second initiation event reads

$$v_{\rm i}^{(2)} = \frac{I(t^+)}{4c_I} = \left[ \frac{N_{\rm B}}{2} \left( 1 + \frac{t^+}{C} \right) + n_{\rm B} \right] / c_{\rm I} = \frac{1}{c_{\rm I}} (\alpha^{(2)} N_{\rm B} + n_{\rm B}). \tag{C5}$$

Hence, the second fixed point reads

$$\alpha^{(2)} = \frac{1}{2} \left( 1 + \frac{t^+}{C} \right), \tag{C6}$$

where $t^+$ is given by Eq. C4. Because of $t^+ < \tau$, this value is smaller than the steady-state $\alpha = (1 + \tau/C)/2$, given by Eq. 3 with $n = 1$. This is consistent with our simulation [Fig. 4(c)]. Thus, the eventual picture is that in the unstable regime, the initiation mass oscillates around the steady-state value Eq. 3 in two values $v_{\rm i}^{(1)}$ and $v_{\rm i}^{(2)}$ given by Eq. C1 and C5.

### Appendix D: Derivation of the steady-state initiation mass formula in the initiator-titration model v2 with a static DnaA-ATP/DnaA-ADP ratio

In our initiator-titration model v2, we first considered a constant DnaA-ATP/DnaA-ADP ratio, named $\gamma$ hereafter, which is likely the case of the $\Delta 4$ mutant [1]. In $\Delta 4$ mutant, there is only DnaA *de novo* synthesis and ATP hydrolysis by intrinsic ATPase activity of DnaA [2]. We denote the intrinsic DnaA-ATP hydrolysis rate as $\nu$. We further assume that this hydrolysis rate is the same for free DnaA-ATP or bound DnaA-ATP on chromosomal binding sites. Additionally, since the free ATP/ADP ratio is high in the cytoplasm, we assume that newly expressed DnaA will immediately form DnaA-ATP. Denoting the total number of DnaA-ATP and DnaA-ADP as $I^{\mathrm{T}}$ and $I^{\mathrm{D}}$, respectively, we have

$$\frac{\mathrm{d}I^{\mathrm{T}}}{\mathrm{d}t} = \lambda(I^{\mathrm{T}} + I^{\mathrm{D}}) - \nu I^{\mathrm{T}}, \tag{D1a}$$

$$\frac{\mathrm{d}I^{\mathrm{D}}}{\mathrm{d}t} = \nu I^{\mathrm{T}}. \tag{D1b}$$

For DnaA-ATP and DnaA-ADP concentrations, $[I^{\mathrm{T}}] = I^{\mathrm{T}}/V$ and $[I^{\mathrm{D}}] = I^{\mathrm{D}}/V$, we have

$$\frac{\mathrm{d}[I^{\mathrm{T}}]}{\mathrm{d}t} = \lambda[I^{\mathrm{D}}] - \nu[I^{\mathrm{T}}], \tag{D2a}$$

$$\frac{\mathrm{d}[I^{\mathrm{D}}]}{\mathrm{d}t} = -\lambda[I^{\mathrm{D}}] + \nu[I^{\mathrm{T}}], \tag{D2b}$$

and the steady-state ratio is given by

$$\frac{[I^{\mathrm{T}}]}{[I^{\mathrm{D}}]} \equiv \gamma = \frac{\lambda}{\nu}. \tag{D3}$$

The time scale for the intrinsic DnaA-ATP hydrolysis is about 15 minutes in wild-type *E. coli* [1]. If the doubling time ranges from 15 minutes to 2.5 hours, then $\gamma$ ranges from 0.1 to 1.

Now we consider that there are bound DnaA on the chromosome and free DnaA in the cytoplasm. We assume that the two forms of DnaA have the same strong binding affinity with chromosomal binding sites, but only DnaA-ATP can bind to DnaA boxes at *ori* binding sites with a relatively weak binding affinity. We denote the number of DnaA-ATP and DnaA-ADP bound to the chromosomal binding sites as $I_{\mathrm{b}}^{\mathrm{T}}$ and $I_{\mathrm{b}}^{\mathrm{D}}$, respectively, the number of free DnaA-ATP and DnaA-ADP in the cytoplasm as $I_{\mathrm{f}}^{\mathrm{T}}$ and $I_{\mathrm{f}}^{\mathrm{D}}$, respectively, and the number of DnaA-ATP at *ori* as $I_{\mathrm{o}}^{\mathrm{T}}$. Based on Eq. D3, we have

$$\frac{I_{\mathrm{b}}^{\mathrm{T}} + I_{\mathrm{f}}^{\mathrm{T}} + I_{\mathrm{o}}^{\mathrm{T}}}{I_{\mathrm{b}}^{\mathrm{D}} + I_{\mathrm{f}}^{\mathrm{D}}} = \gamma. \tag{D4}$$

The DnaA binding and unbinding processes are fast compared to the doubling time, so we can assume binding reactions are in rapid equilibrium, which gives

$$I_{\mathrm{f}}^{\mathrm{T}}(B_{\mathrm{tot}} - I_{\mathrm{b}}^{\mathrm{T}} - I_{\mathrm{b}}^{\mathrm{D}}) = I_{\mathrm{b}}^{\mathrm{T}} K_{\mathrm{b}} V, \tag{D5}$$

$$I_{\mathrm{f}}^{\mathrm{D}}(B_{\mathrm{tot}} - I_{\mathrm{b}}^{\mathrm{T}} - I_{\mathrm{b}}^{\mathrm{D}}) = I_{\mathrm{b}}^{\mathrm{D}} K_{\mathrm{b}} V, \tag{D6}$$

$$I_{\mathrm{f}}^{\mathrm{T}}(O_{\mathrm{tot}} - I_{\mathrm{o}}^{\mathrm{T}}) = I_{\mathrm{o}}^{\mathrm{T}} K_{\mathrm{o}} V, \tag{D7}$$

where $K_{\mathrm{b}}$ and $K_{\mathrm{o}}$ are the dissociation constant for chromosomal binding sites and for *ori* binding sites, respectively; $B_{\mathrm{tot}}$ is the total number of chromosomal binding sites, and $O_{\mathrm{tot}}$ is the total *ori* binding sites that must be larger than $n_{\mathrm{B}}$ at each *ori*. Moreover, we have the equation of balanced biosynthesis for total DnaA,

$$c_{\mathrm{I}} V = I_{\mathrm{b}}^{\mathrm{T}} + I_{\mathrm{f}}^{\mathrm{T}} + I_{\mathrm{o}}^{\mathrm{T}} + I_{\mathrm{b}}^{\mathrm{D}} + I_{\mathrm{f}}^{\mathrm{D}}. \tag{D8}$$

To determine the initiation mass $v_{\mathrm{i}}$, suppose there are $2^d$ *ori*'s at initiation, then $V = 2^d v_{\mathrm{i}}$, $I_{\mathrm{o}}^{\mathrm{T}} = 2^d n_{\mathrm{B}}$, and $O_{\mathrm{tot}} - I_{\mathrm{o}}^{\mathrm{T}} = 2^d \Delta_{\mathrm{B}}$, where $\Delta_{\mathrm{B}}$ is the number of remaining binding sites at each *ori* at initiation. As discussed in Appendix A and B, $B_{\mathrm{tot}}$ at initiation is fully determined by the replication fork progress that is a function of growth rates: $B_{\mathrm{tot}} = 2^d \alpha N_{\mathrm{B}}$, where $\alpha$ was defined in Eq. 3 in the main text.

In principle, by substituting these parameters into Eqs. D4 - D8, we can solve the unknown variables $I_{\mathrm{f}}^{\mathrm{T}}$, $I_{\mathrm{f}}^{\mathrm{D}}$, $I_{\mathrm{b}}^{\mathrm{T}}$, $I_{\mathrm{b}}^{\mathrm{D}}$, and $v_{\mathrm{i}}$. However, it is hard to obtain a simple expression by solving Eq. D4 - D8 directly because we need to solve

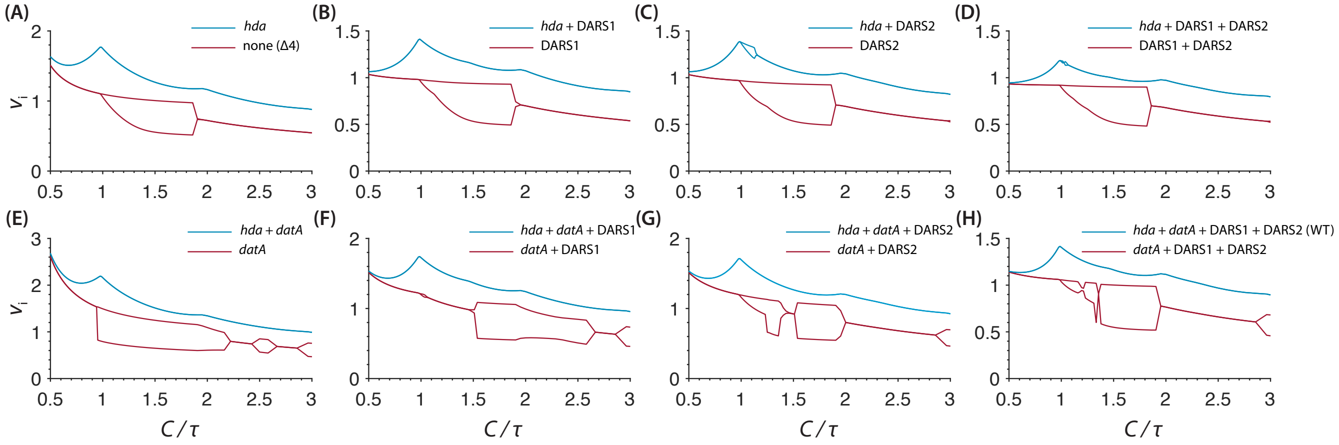


FIG. 3. The initiation mass behavior in different combinations of the DnaA-ATP/DnaA-ADP conversion elements, i.e., *hda* (RIDA), *datA* (DDAH), DARS1, and DARS2. The situations with *hda* and without *hda* are plotted together in each plots to show the contrast.

a polynomial equation. To simplify the solution, notice that $B_{\rm tot}$ should be almost saturated at initiation because of strong binding affinity, so approximately

$$I_{\rm b}^{\rm T} + I_{\rm b}^{\rm D} \approx B_{\rm tot}. \tag{D9}$$

On the other hand, since $0.1 \leq \gamma \leq 1$, $I_{\rm b}^{\rm T} \sim \gamma/(1+\gamma)B_{\rm tot} > I_{\rm o}^{\rm T}$, so we can fairly assume $I_{\rm b}^{\rm T} + I_{\rm f}^{\rm T} \gg I_{\rm o}^{\rm T}$. By Eqs. D5 and D6, Eq. D4 becomes

$$\frac{I_{\rm b}^{\rm T}}{I_{\rm b}^{\rm D}} \approx \frac{I_{\rm f}^{\rm T} + I_{\rm o}^{\rm T}}{I_{\rm f}^{\rm D}} = \gamma. \tag{D10}$$

These approximations make it easy to solve the original Eqs. D4 - D8, and finally we obtain

$$v_{\rm i} = \frac{\alpha N_{\rm B} + (1+\frac{1}{\gamma})n_{\rm B}}{c_{\rm I} - (1+\frac{1}{\gamma})K_{\rm eff} n_{\rm B}}, \tag{D11}$$

where $K_{\rm eff} = K_{\rm o}/\Delta_{\rm B}$. This is exactly Eq. 9 in the main text.

The dissociation constant $K_{\rm o}$ at *ori* should be much larger than $K_{\rm b}$ in most of the time. In our simulation, we set $c_{\rm I} = 400\ \mu{\rm m}^{-3}$, $n_{\rm B} = 1$, $K_{\rm b} = 1\ \mu{\rm m}^{-3}$, and $K_{\rm o} > 10\ \mu{\rm m}^{-3}$ [3]. These values result in a very large $v_{\rm i}$. This is because we do not assume cooperativity of DnaA binding to *ori*, so that the binding process at *ori* will be slowed down significantly when the occupancy is close to the threshold $n_{\rm B}$. To resolve this effect, we consider that $K_{\rm o}$ can decrease with $I_{\rm o}^{\rm T}$ because of stabilized filamentous structure of DnaA proteins at *ori*, which has been found in *E. coli* [4]. In this case, the rapid-equilibrium assumption may not hold, but we assume that it is not too far away. Hence, we treat $K_{\rm eff}$ as a fitting parameter. In our simulation, we assumed a linear decrease of $K_{\rm o}$ from 50 $\mu{\rm m}^{-3}$ to 1 $\mu{\rm m}^{-3}$ when the occupancy rises from 0 to the threshold. We fitted the numerical results with the initiation expression Eq. D11, and found that $K_{\rm eff} = 2\ \mu{\rm m}^{-3}$ is almost a perfect fit for the stable regimes [Fig. 5(c) in the main text].

### Appendix E: The numerical model set-up and supplementary results

In this section, we include the effects of RIDA, DDAH, DARS1 and DARS2. First, it is reported that RIDA accelerates the conversion of DnaA-ATP to DnaA-ADP by the Hda protein that binds to the DNA-loaded $\beta$-clamp at the replisome [5–7]. We assume that Hda proteins work at the saturation level, so the RIDA activity is proportional to the number of replication forks, $N_{\rm fork}$. Second, DDAH promotes the conversion of DnaA-ATP to DnaA-ADP by the locus *datA* on the chromosome [8, 9]. Thus, we assume DDAH activity to be proportional to the copy number of *datA* loci, $N_{\rm datA}$. Third, DARS1 and DARS2 are the two loci on the chromosome that promotes the reactivation

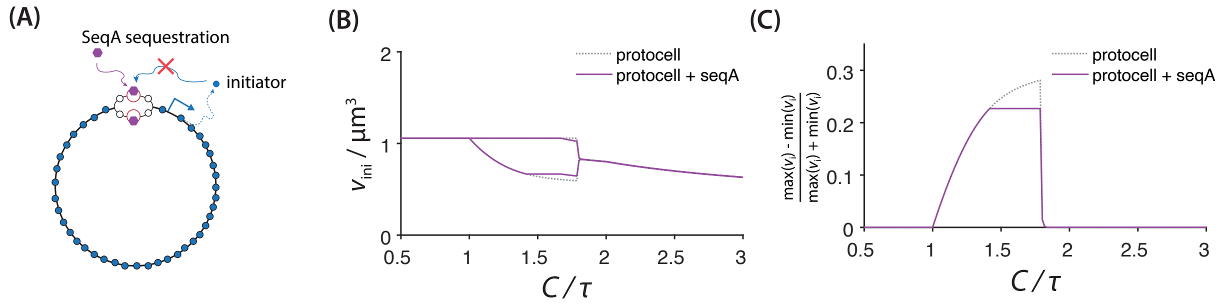


FIG. 4. The effect of SeqA in the protocell model. (a) We consider the SeqA sequestration effect at *ori* binding sites, which results in no initiation events in a duration time of $t_{\rm seq} = 10$ min. (b) The unstable range does not shrink with the existence of seqA. (c) The amplitude of the initiation mass oscillation, defined by $(\max v_{\rm i} - \min v_{\rm i})/(\max v_{\rm i} + \min v_{\rm i})$, is slightly reduced when $C/\tau$ is relatively large in the unstable regime. Here, we set $n_B/N_B = 1/30$.

of DnaA-ADP [10, 11], hence, we assume that their activities are proportional to the copy number of DARS1 and DARS2 loci, $N_{\rm DARS1}$ and $N_{\rm DARS2}$, respectively. We further assume that these reactions follow the Michaelis-Menten form. Based on these assumptions, we have

$$\frac{{\rm d}I^{\rm T}}{{\rm d}t} = \lambda(I^{\rm T}+I^{\rm D}) - \nu I^{\rm T} - (k_{\rm RIDA}N_{\rm fork} + k_{\rm DDAH}N_{\rm datA})\frac{[I^{\rm T}]}{[I^{\rm T}]+K_{\rm T}} + (k_{\rm DARS1}N_{\rm DARS1} + k_{\rm DARS2}N_{\rm DARS2})\frac{[I^{\rm D}]}{[I^{\rm D}]+K_{\rm D}}, \tag{E1a}$$

$$\frac{{\rm d}I^{\rm D}}{{\rm d}t} = \nu I^{\rm T} + (k_{\rm RIDA}N_{\rm fork} + k_{\rm DDAH}N_{\rm datA})\frac{[I^{\rm T}]}{[I^{\rm T}]+K_{\rm T}} - (k_{\rm DARS1}N_{\rm DARS1} + k_{\rm DARS2}N_{\rm DARS2})\frac{[I^{\rm D}]}{[I^{\rm D}]+K_{\rm D}}, \tag{E1b}$$

where $k_{\rm RIDA}$, $k_{\rm DDAH}$, $k_{\rm DARS1}$, and $k_{\rm DARS2}$ are the reaction rate constants corresponding to RIDA, DDAH, DARS1, and DARS2, respectively; $K_{\rm T}$ and $K_{\rm D}$ are the Michaelis-Menten constants for DnaA-ATP to DnaA-ADP conversion and the other way around, respectively. We have assumed that the Michaelis-Menten constant is the same for RIDA and DDAH, and for DARS1 and DARS2, for simplicity. When we study the absence of some of the mechanisms, we set the corresponding reaction rate constants to be zero.

We are particularly interested in the dynamics of $\gamma = I^{\rm T}/I^{\rm D}$. Based on Eqs. E1, we can obtain the ODE for $\gamma$:

$$\frac{{\rm d}\gamma}{{\rm d}t} = (1+\gamma)\left[\lambda - \nu\gamma - \frac{k_{\rm RIDA}N_{\rm fork} + k_{\rm DDAH}N_{\rm datA}}{c_{\rm I}V}\frac{\gamma}{\frac{\gamma}{1+\gamma}+K_{\rm T}^*} + \frac{k_{\rm DARS1}N_{\rm DARS1} + k_{\rm DARS2}N_{\rm DARS2}}{c_{\rm I}V}\frac{1}{\frac{1}{1+\gamma}+K_{\rm D}^*}\right], \tag{E2}$$

where $K_{\rm T}^* = K_{\rm T}/c_{\rm I}$ and $K_{\rm D}^* = K_{\rm D}/c_{\rm I}$.

Now we consider the bound and unbound forms of DnaA: DnaA bound to chromosomal binding sites $I_{\rm b} = I_{\rm b}^{\rm T} + I_{\rm b}^{\rm D}$, free DnaA in cytoplasm $I_{\rm f} = I_{\rm f}^{\rm T} + I_{\rm f}^{\rm D}$, and DnaA bound to *ori* binding sites $I_{\rm o}$. Eq. D10 is still assumed here. We denote the binding and unbinding rate constant on the chromosomal binding sites as $k_{\rm on}^{\rm b}$ and $k_{\rm off}^{\rm b}$, and those on the *ori* binding sites as $k_{\rm on}^{\rm o}$ and $k_{\rm off}^{\rm o}$. Thus, The ODEs for $I_{\rm b}$, $I_{\rm f}$, and $I_{\rm o}$ read

$$\frac{{\rm d}I_{\rm b}}{{\rm d}t} = k_{\rm on}^{\rm b}\frac{I_{\rm f}}{V}(B_{\rm tot} - I_{\rm b}) - k_{\rm off}^{\rm b}I_{\rm b}, \tag{E3a}$$

$$\frac{{\rm d}I_{\rm f}}{{\rm d}t} = \lambda(I_{\rm f}+I_{\rm b}+I_{\rm o}) - k_{\rm on}^{\rm b}\frac{I_{\rm f}}{V}(B_{\rm tot} - I_{\rm b}) + k_{\rm off}^{\rm b}I_{\rm b} - \frac{\gamma}{1+\gamma}k_{\rm on}^{\rm o}\frac{I_{\rm f}}{V}(O_{\rm tot} - I_{\rm o}) + k_{\rm off}^{\rm o}I_{\rm o} \tag{E3b}$$

$$\frac{{\rm d}I_{\rm o}}{{\rm d}t} = \frac{\gamma}{1+\gamma}k_{\rm on}^{\rm o}\frac{I_{\rm f}}{V}(O_{\rm tot} - I_{\rm o}) - k_{\rm off}^{\rm o}I_{\rm o} \tag{E3c}$$

where $B_{\rm tot}$ follows the same dynamics determined by $\boldsymbol{\rho}(t)$ as derived in Appendix B. In this way, we conducted deterministic simulation based on Eq. E2 and Eqs. E3. The results with all the sixteen combinations of the four mechanisms are in Fig. 3.

#### Appendix F: Relation between the intrinsic/extrinsic noise and initiation asynchrony/cell-to-cell variability

We defined the intrinsic noise and the extrinsic noise for the initiation mass based on one of their conventional definitions [12] in Section E. However, we will show they are not equivalent to the intuitive concepts of asynchrony and cell-to-cell variability, though these two groups of concepts are closely related.

For asynchrony vs. cell-to-cell variability, by intuition, we can quantify them as the coefficients of variation along the diagnal axis and off-diagnal axis on the ori1 - ori2 initiation mass plot [Fig. 6(a) in the main text], respectively. That is, if we transform the initiation mass variables $v_{\rm i}^{(1)}$ and $v_{\rm i}^{(2)}$ by a rotation of the cooordinates, $u_{\rm i}^{(1)} = (v_{\rm i}^{(1)} + v_{\rm i}^{(2)})/\sqrt{2}$, $u_{\rm i}^{(2)} = (v_{\rm i}^{(1)} - v_{\rm i}^{(2)})/\sqrt{2}$, initiation cell-to-cell variability ($CV_{\rm ccv}$) and initiation asynchrony ($CV_{\rm asyn}$) can be properly defined as

$$CV_{\rm ccv}^2 = \frac{\left\langle u_{\rm i}^{(1)\ 2} \right\rangle - \left\langle u_{\rm i}^{(1)} \right\rangle^2}{\left\langle u_{\rm i}^{(1)} \right\rangle^2}, \quad CV_{\rm asyn}^2 = \frac{\left\langle u_{\rm i}^{(2)\ 2} \right\rangle - \left\langle u_{\rm i}^{(2)} \right\rangle^2}{\left\langle u_{\rm i}^{(1)} \right\rangle^2}. \tag{F1}$$

We want to use the statistics of $v_{\rm i}^{(1)}$ and $v_{\rm i}^{(2)}$ to represent these two new CVs. We assume that $\left\langle v_{\rm i}^{(1)} \right\rangle = \left\langle v_{\rm i}^{(2)} \right\rangle$, and $\left\langle v_{\rm i}^{(1)\ 2} \right\rangle = \left\langle v_{\rm i}^{(2)\ 2} \right\rangle$, because two *ori*'s are identical such that $v_{\rm i}^{(1)}$ and $v_{\rm i}^{(2)}$ should follow the same distribution. Therefore, their coefficients of variation should be the same and equal to the total CV:

$$CV^{(1)\ 2} = CV^{(2)\ 2} = CV_{\rm tot}^2 = \frac{\langle v_{\rm i}^2 \rangle - \langle v_{\rm i} \rangle^2}{\langle v_{\rm i} \rangle^2}. \tag{F2}$$

Substituting the definition of $u_{\rm i}^{(1)}$ and $u_{\rm i}^{(2)}$ to Eq. F1, we have

$$CV_{\rm ccv}^2 = \frac{1+r}{2} CV_{\rm tot}^2, \quad CV_{\rm asyn}^2 = \frac{1-r}{2} CV_{\rm tot}^2, \tag{F3}$$

where $r$ is the Pearson correlation coefficient between $v_{\rm i}^{(1)}$ and $v_{\rm i}^{(2)}$,

$$r = \frac{\left\langle v_{\rm i}^{(1)} v_{\rm i}^{(2)} \right\rangle - \left\langle v_{\rm i}^{(1)} \right\rangle \left\langle v_{\rm i}^{(2)} \right\rangle}{\sqrt{\left\langle v_{\rm i}^{(1)\ 2} \right\rangle - \left\langle v_{\rm i}^{(1)} \right\rangle^2} \cdot \sqrt{\left\langle v_{\rm i}^{(2)\ 2} \right\rangle - \left\langle v_{\rm i}^{(2)} \right\rangle^2}}. \tag{F4}$$

On the other hand, based on the conventional definition [12], the intrinsic noise and the extrinsic noise for the initiation mass are defined in Eq. 12 in Section E1. Following the above derivations, we have

$$CV_{\rm int}^2 = (1-r) CV_{\rm tot}^2, \quad CV_{\rm ext}^2 = r CV_{\rm tot}^2. \tag{F5}$$

However, this definition is only valid when $0 \le r \le 1$ given that $CV_{\rm ext}$ has to be a real number, which means it only applies when two variables are non-negatively correlated. Based on Eqs. F3 & F5, we obtain the relation between intrinsic/extrinsic noise and asynchrony/cell-to-cell variability,

$$CV_{\rm asyn}^2 = \frac{1}{2} CV_{\rm int}^2, \quad CV_{\rm ccv}^2 = CV_{\rm ext}^2 + \frac{1}{2} CV_{\rm int}^2. \tag{F6}$$

We have the summation relation of the two groups of concepts,

$$CV_{\rm ccv}^2 + CV_{\rm asyn}^2 = CV_{\rm ext}^2 + CV_{\rm int}^2 = CV_{\rm tot}^2. \tag{F7}$$

Hence, asynchrony/cell-to-cell variability and the intrinsic/extrinsic noise are just two decompositions of the total noise. The first works only at the regime $0 \le r \le 1$, while the second works for any correlation situation, i.e., $-1 \le r \le 1$.

#### Appendix G: Derivation of the CVs' scaling laws in the first-passage-time (FPT) models

Our first model in Section E2 assumes that 1. the initiator proteins are produced in a Poisson process with a constant production rate $\beta$; 2. after production, the initiator protein has equal chance to bind to either *ori*1 or *ori*2, namely, a Bernoulli trial. Thus, the probability of *ori*1 to accumulate one protein during the time interval $(t, t+\mathrm{d}t]$ reads

$$\mathbb{P}[O_1(t+\mathrm{d}t) = m+1 | O_1(t) = m] = \beta \mathrm{d}t \cdot \frac{1}{2}, \tag{G1}$$

which is equivalently a Poisson process with a production rate of $\beta' = \frac{\beta}{2}$.

For a Poisson process with a production rate of $\beta'$, the waiting time for each jump event follows a exponential distribution with a decay rate $\beta'$, and the first-passage-time (FPT) $T^{(1)}$ is the sum of $n_B$ identical waiting-time variables. This gives out a Gamma distribution of $T^{(1)}$,

$$P(T; n_\mathrm{B}, \beta') = \frac{\beta^{n_\mathrm{B}} T^{n_\mathrm{B}-1} e^{-\beta' T}}{(n_\mathrm{B}-1)!}, \tag{G2}$$

Thus, $\left\langle T^{(1)} \right\rangle = n_\mathrm{B}/\beta' = 2n_\mathrm{B}/\beta$, and $\sigma_{T^{(1)}} = \sqrt{n_\mathrm{B}}/\beta' = 2\sqrt{n_\mathrm{B}}/\beta$. Based on Eq. 14, we obtain

$$CV_\mathrm{int} = \frac{1}{\sqrt{n_\mathrm{B}}}. \tag{G3}$$

On the other hand, the total number of proteins produced before time $T$ follows the original Poisson distribution,

$$P(N; T) = \frac{(\beta T)^N}{N!} e^{-\beta T}, \tag{G4}$$

which gives a mean value $\langle N \rangle = \beta T$. Thus, the mean total protein at the FPT ($\left\langle T^{(1)} \right\rangle = 2n_\mathrm{B}/\beta$) is $\langle N \rangle = 2n_\mathrm{B}$. Therefore,

$$CV_\mathrm{int} = \sqrt{\frac{2}{\langle N \rangle}}, \tag{G5}$$

which is essentially Eq. 15.

Next, for the two-step Poisson process, we have decomposed $T^{(1)}$ and $T^{(2)}$ into three independent variables, $T^{(1)} = T^{(0)} + \Delta T^{(1)}$, $T^{(2)} = T^{(0)} + \Delta T^{(2)}$ in the main text. To calculate Eq. 16, we need to obtain $\left\langle T^{(1)} \right\rangle$, $\sigma_{\Delta T^{(1)}}$, and $\sigma_{T^{(0)}}$. To avoid the instability issue, we consider the stable cases of two-overlapping cell cycle (non-multifork replication). In this situation, The threshold for the titration step is roughly $2N_\mathrm{B}$, and for the *ori*1 accumulation step is $n_B$. The first titration step has an accumulation rate of $\beta$, while the second step has an accumulation rate of $\beta' = \beta/2$, similar to the simple Poisson process case above. Thus,

$$\left\langle T^{(0)} \right\rangle = \frac{2N_\mathrm{B}}{\beta}, \quad \left\langle \Delta T^{(1)} \right\rangle = \frac{n_B}{\beta'} = \frac{2n_B}{\beta}; \tag{G6}$$

$$\sigma_{T^{(0)}} = \frac{\sqrt{2N_\mathrm{B}}}{\beta}, \quad \sigma_{\Delta T^{(1)}} = \frac{\sqrt{n_\mathrm{B}}}{\beta'} = \frac{2\sqrt{n_\mathrm{B}}}{\beta}. \tag{G7}$$

Hence, $\langle N \rangle = \beta \left\langle T^{(1)} \right\rangle = 2(N_\mathrm{B} + n_\mathrm{B})/\beta$. We rewrite Eq. G7 in terms of $\langle N \rangle$ and obtain

$$CV_\mathrm{int} = \frac{2\sqrt{n_\mathrm{B}}}{\langle N \rangle}, \quad CV_\mathrm{ext} = \frac{\sqrt{\langle N \rangle - 2n_\mathrm{B}}}{\langle N \rangle} \approx \frac{1}{\sqrt{\langle N \rangle}}, \tag{G8}$$

which is the same as Eq. 17. in the main text.

To verify that the decomposition is valid, we performed Monte Carlo simulation. In simulation, the initiator protein is generated based on the Poisson process with a constant rate $\beta$. Once a protein is generated, it will undergo a trial among titration sites, *ori*1, and *ori*2. The probability of the protein to bind to different destinations is proportional to their binding affinities, and we set the binding affinity difference between titration sites and *ori*'s to be 100 fold, based on the *E. coli* case we mentioned in the main text. Further, we assumed that when the occupancy of one *ori* reaches the threshold, it will keep accumulation rather than waiting for the other *ori*. This is to assure that the stochastic process is not changed after one *ori* reaches the threshold. We notice that although these detailed settings are necessary to get numbers very close to what are predicted in Eq. 17 [Fig. 6(d), dashed grey lines], the $1/N$ vs. $1/\sqrt{N}$ scaling laws are quite conserved in settings with different details.

[1] T. Boesen, G. Charbon, H. Fu, C. Jensen, D. Li, S. Jun, and others, Robust control of replication initiation in the absence of DnaA-ATP DnaA-ADP regulatory elements in escherichia coli, bioRxiv (2022).
[2] K. Sekimizu, D. Bramhill, and A. Kornberg, ATP activates dnaa protein in initiating replication of plasmids bearing the origin of the e. coli chromosome, Cell **50**, 259 (1987).
[3] S. Schaper and W. Messer, Interaction of the initiator protein DnaA of escherichia coli with its DNA target, J. Biol. Chem. **270**, 17622 (1995).
[4] T. Katayama, K. Kasho, and H. Kawakami, The DnaA cycle in escherichia coli: Activation, function and inactivation of the initiator protein, Front. Microbiol. **8**, 2496 (2017).
[5] T. Katayama, T. Kubota, K. Kurokawa, E. Crooke, and K. Sekimizu, The initiator function of DnaA protein is negatively regulated by the sliding clamp of the e. coli chromosomal replicase, Cell **94**, 61 (1998).
[6] J. Kato and T. Katayama, Hda, a novel DnaA-related protein, regulates the replication cycle in escherichia coli, EMBO J. **20**, 4253 (2001).
[7] M. C. Moolman, S. T. Krishnan, J. W. J. Kerssemakers, A. van den Berg, P. Tulinski, M. Depken, R. Reyes-Lamothe, D. J. Sherratt, and N. H. Dekker, Slow unloading leads to DNA-bound $\beta$2-sliding clamp accumulation in live escherichia coli cells, Nat. Commun. **5**, 5820 (2014).
[8] R. Kitagawa, H. Mitsuki, T. Okazaki, and T. Ogawa, A novel DnaA protein-binding site at 94.7 min on the escherichia coli chromosome, Mol. Microbiol. **19**, 1137 (1996).
[9] K. Kasho and T. Katayama, DnaA binding locus *datA* promotes DnaA-ATP hydrolysis to enable cell cycle-coordinated replication initiation, Proceedings of the National Academy of Sciences **110**, 936 (2013).
[10] K. Fujimitsu, T. Senriuchi, and T. Katayama, Specific genomic sequences of e. coli promote replicational initiation by directly reactivating ADP-DnaA, Genes Dev. **23**, 1221 (2009).
[11] K. Kasho, K. Fujimitsu, T. Matoba, T. Oshima, and T. Katayama, Timely binding of IHF and fis to DARS2 regulates ATP-DnaA production and replication initiation, Nucleic Acids Res. **42**, 13134 (2014).
[12] M. B. Elowitz, A. J. Levine, E. D. Siggia, and P. S. Swain, Stochastic gene expression in a single cell, Science **297**, 1183 (2002).
